# Machine learning improves global models of plant diversity

**DOI:** 10.1101/2022.04.08.487610

**Authors:** Lirong Cai, Holger Kreft, Amanda Taylor, Pierre Denelle, Julian Schrader, Franz Essl, Mark van Kleunen, Jan Pergl, Petr Pyšek, Anke Stein, Marten Winter, Julie F. Barcelona, Nicol Fuentes, Inderjit, Dirk Nikolaus Karger, John Kartesz, Andreij Kuprijanov, Misako Nishino, Daniel Nickrent, Arkadiusz Nowak, Annette Patzelt, Pieter B. Pelser, Paramjit Singh, Jan J. Wieringa, Patrick Weigelt

## Abstract

Despite the paramount role of plant diversity for ecosystem functioning, biogeochemical cycles, and human welfare, knowledge of its global distribution is incomplete, hampering basic research and biodiversity conservation. Here, we used machine learning (random forests, extreme gradient boosting, neural networks) and conventional statistical methods (generalised linear models, generalised additive models) to model species richness and phylogenetic richness of vascular plants worldwide based on 830 regional plant inventories including c. 300,000 species and predictors of past and present environmental conditions. Machine learning showed an outstanding performance, explaining up to 80.9% of species richness and 83.3% of phylogenetic richness. Current climate and environmental heterogeneity emerged as the primary drivers, while past environmental conditions left only small but detectable imprints on plant diversity. Finally, we combined predictions from multiple modelling techniques (ensemble predictions) to reveal global patterns and centres of plant diversity at multiple resolutions down to 7,774 km^2^. Our predictive maps provide the most accurate estimates of global plant diversity available to date at grain sizes relevant for conservation and macroecology.

## Introduction

Vascular plants comprise well over 340,000 species^1^ and are fundamental to terrestrial ecosystems maintaining ecosystem functioning^2^ and providing ecosystem services^3,4^. To preserve and manage this important part of global biodiversity, knowledge of its spatial distribution is critical. Mapping of plant distributions and diversity can be traced back to Alexander von Humboldt in the 19^th^ century and subsequent pioneering scientific contributions (reviewed in Mutke and Barthlott^5^). However, even today, we still lack a detailed understanding of the distribution of all vascular plant species worldwide. We, therefore, rely on modelling techniques to overcome these shortcomings and to predict diversity patterns in relation to environmental and spatial variables^6^. The accuracy of such predictive maps depends on the quality and representativeness of the available plant diversity data, environmental predictors, and the modelling techniques applied.

Knowledge of plant distributions worldwide has increased tremendously in recent years, thanks to immense international efforts to mobilize and collate species occurrence records^7,8^, vegetation plots^9^ and regional checklists and Floras^1,10^. However, these data differ in precision, completeness, and scope^11^. Specifically, fine-grained data such as occurrence records and vegetation plots are often geographically biased and only partially cover regional floras^12,13^. Despite being coarse-grained and often delimited by artificial, administrative borders, checklists and Floras reflect fairly complete and authoritative accounts of the floristic composition of differently sized regions, and are available with near-complete global coverage^1,10^. As such, checklists and Floras are immensely useful resources for global-scale modelling of plant diversity–environment relationships^6^, and for predicting plant diversity across different grain sizes^14^. The inclusion of species identities further allows for the integration of species-level phylogenetic and trait information, offering a unique opportunity to study multiple facets of biodiversity.

Several hypotheses have been proposed to explain global plant diversity patterns^15–17^. Although it is widely accepted that plant diversity reflects a complex interplay between evolutionary, geological, and ecological processes, disentangling the different drivers of plant diversity remains at the forefront of modern macroecology^6,18^. Large and heterogeneous areas, for example, may support more species by offering a greater diversity of resources and habitats, thus promoting species coexistence^19^ and offering refugia during environmental fluctuations^20^. Areas with warm, wet and relatively stable climates such as humid tropical regions may also support more species owing to high speciation^17,21,22^ and low extinction rates^23,24^. Geographic isolation may simultaneously promote extinction^25,26^ and speciation^27^, by making populations less well-connected. Finally, historical processes like plate tectonics and climatic change have influenced diversity patterns through altered biotic isolation and exchange or species range shifts^28–30^. However, past environmental conditions remain underrepresented in global models of plant diversity and their legacies in modern plant distributions are still poorly understood^31,32^.

Diversity–environment relationships are often complex, non-linear, and scale-dependent^14,33^. Many environmental predictors interact and show high levels of collinearity, thus presenting major challenges for conventional statistical models such as generalised linear models (GLMs) and generalised additive models (GAMs). Machine learning approaches represent powerful modelling tools that can effectively deal with multidimensional and correlated data and can reveal non-linear relationships and interactions of predictors without *a priori* specification^34,35^. Therefore, machine learning has become a promising alternative to conventional techniques in ecology^36–39^. However, its performance in modelling global plant diversity has yet to be explored. In addition to relying on one particular model type, combining predictions based on multiple modelling techniques (i.e. ensemble predictions) may decrease prediction uncertainties^40^ and can thereby further improve predictions of global plant diversity patterns.

Here, we present improved models and predictions of two key facets of vascular plant diversity, i.e. species richness and phylogenetic richness (measured as Faith’s PD^41^), at a global extent using advanced statistical modelling techniques. In addition to non-spatial and spatial GLMs and GAMs, we systematically assess the predictive performance of machine learning methods, including random forests, extreme gradient boosting (XGBoost) and neural networks. Specifically, our aims are: (1) to compare the performance of different modelling techniques in revealing complex diversity–environment relationships and to improve global geo-statistical plant diversity models; (2) to quantify the relative importance of past and current environmental drivers of plant species richness and phylogenetic richness; and (3) to predict both facets of plant diversity at multiple grain sizes across the globe. Our study is based on nearly 300,000 species from checklists and Floras for 830 regions across the globe collated in the Global Inventory of Floras and Traits^10^ (GIFT; http://gift.uni-goettingen.de; Supplementary references 2), and a large, dated mega-phylogeny of vascular plants^42^ (Supplementary Fig. 1).

## Results

### Performance of plant diversity models

Our results reveal a great potential of machine learning, particularly decision-tree methods, for modelling plant diversity–environment relationships and accurately predicting plant diversity across various scales. We generated models for species richness and phylogenetic richness using five different modelling techniques (Table 1) and 15 predictor variables representing geography, climate, environmental heterogeneity and past environments (Supplementary Table 1). Predictors were selected to represent the major hypotheses related to plant diversity–environment relationships^6,14,20,32,33^ and were filtered based on their contribution to model performance and collinearity (Supplementary Table 1). Overall, the predictive power of the models was high (Table 1). Machine learning models and GAMs outperformed GLMs, and spatial models (i.e. models containing spatial terms to account for the spatial non-independence of regions^43^) showed an overall better performance than non-spatial models (except GLMs for species richness). Extreme gradient boosting, an ensemble of sequentially trained decision trees, produced the most accurate predictions for both species richness (70.3% variation explained based on spatial cross-validation, 80.9% based on random cross-validation) and phylogenetic richness (73.7 and 83.3%, respectively), which was consistent across spatial and non-spatial models.

**Table 1.** Performance of global models of vascular plant diversity based on cross validation.

| Models | Species richness |  |  |  | Phylogenetic richness (Faith's PD) |  |  |  |
| --- | --- | --- | --- | --- | --- | --- | --- | --- |
|  | Random cross-validation |  | Spatial cross-validation |  | Random cross-validation |  | Spatial cross-validation |  |
|  | RMSE | R<sup>2</sup> | RMSE | R<sup>2</sup> | RMSE | R<sup>2</sup> | RMSE | R<sup>2</sup> |
| **Non-spatial models** |  |  |  |  |  |  |  |  |
| Full GLM | 0.525 | 0.636 | 0.582 | 0.561 | 0.514 | 0.452 | 0.552 | 0.359 |
| Minimum adequate GLM | 0.520 | 0.643 | 0.548 | 0.608 | 0.513 | 0.454 | 0.548 | 0.369 |
| GLM with interaction terms | 0.471 | 0.704 | 0.502 | 0.664 | 0.412 | 0.635 | 0.453 | 0.559 |
| GAM | 0.437 | 0.742 | 0.507 | 0.658 | 0.359 | 0.723 | 0.430 | 0.604 |
| Random forests | 0.415 | 0.761 | 0.511 | 0.639 | 0.317 | 0.784 | 0.395 | 0.667 |
| Extreme gradient boosting | 0.389 | 0.791 | 0.487 | 0.673 | 0.295 | 0.813 | 0.384 | 0.685 |
| Neural networks | 0.451 | 0.718 | 0.604 | 0.496 | 0.328 | 0.769 | 0.419 | 0.628 |
| **Spatial models** |  |  |  |  |  |  |  |  |
| SAR | 0.537 | 0.600 | 0.548 | 0.584 | 0.416 | 0.629 | 0.426 | 0.611 |
| GAM | 0.413 | 0.769 | 0.499 | 0.667 | 0.340 | 0.751 | 0.416 | 0.633 |
| Random forests | 0.398 | 0.780 | 0.502 | 0.653 | 0.303 | 0.803 | 0.379 | 0.694 |
| Extreme gradient boosting | 0.371 | 0.809 | 0.463 | 0.703 | 0.279 | 0.833 | 0.351 | 0.737 |
| Neural networks | 0.422 | 0.753 | 0.587 | 0.522 | 0.314 | 0.789 | 0.433 | 0.597 |

Each model was evaluated for its predictive performance using both random 10-fold and spatial 68-fold cross-validation. Non-spatial models were fitted with 15 predictors representing geography, climate, environmental

The good predictive performance of machine learning models can be attributed to their ability to uncover complex, non-linear diversity–environment relationships (Supplementary Figs. 2 and 3) and interactive effects (Supplementary Figs. 4-15). We found strong interactions between spatial terms and environmental variables (Supplementary Figs. 4-15). This indicates regional differences in plant diversity and diversity–environment relationships and shows that different combinations of environmental variables are important when predicting diversity across geographic regions^14^. Moreover, machine learning models revealed strong interactions between energy and water availability, energy and environmental heterogeneity, as well as area and environmental variables (Supplementary Figs. 4-15). Also, the accuracy of GLM increased when including the interactions that turned out to be important in machine learning models (70.4% vs. 63.6% in species richness based on random cross-validation; 63.5% vs. 45.2% in phylogenetic richness), highlighting the role of complex interactive effects among biotic and abiotic factors in shaping global plant diversity patterns^6,14,33^. By implicitly accounting for grain dependence and complex interactions among spatial and environmental variables, our machine learning models outperform previous models of plant diversity^6,14^ (Supplementary Table 2), improving our understanding of diversity–environment relationships and allowing for improved predictions of plant diversity across scales.

### Drivers of global patterns of vascular plant diversity

Current climatic variables emerged as the most important drivers of plant diversity, accounting for 34.4-48.1% of the variation in species richness and 39.7-58.2% in phylogenetic richness across models (Fig. 1; Supplementary Table 1). High energy and water availability and low seasonality promoted species and phylogenetic richness (Supplementary Figs. 2 and 3), supporting other large-scale studies that report strong effects of current climate on plant diversity^6,33,44^. Environmental heterogeneity (measured here as elevational range and number of soil types within a region) explained 21.0-40.9% of the variation in species richness and 16.3-27.2% in phylogenetic richness, with increasing heterogeneity leading to higher plant diversity as expected^20^. Even though species and phylogenetic richness were highly correlated (Pearson’s r = 0.98), some differences emerged in diversity–environment relationships. For example, environmental heterogeneity explained less variation in phylogenetic richness than in species richness. This potentially reflects a signal of *in-situ* speciation that is promoted by high environmental heterogeneity, creating clusters of closely related species resulting in relatively low phylogenetic richness compared to species richness^45^.

**Fig. 1.**
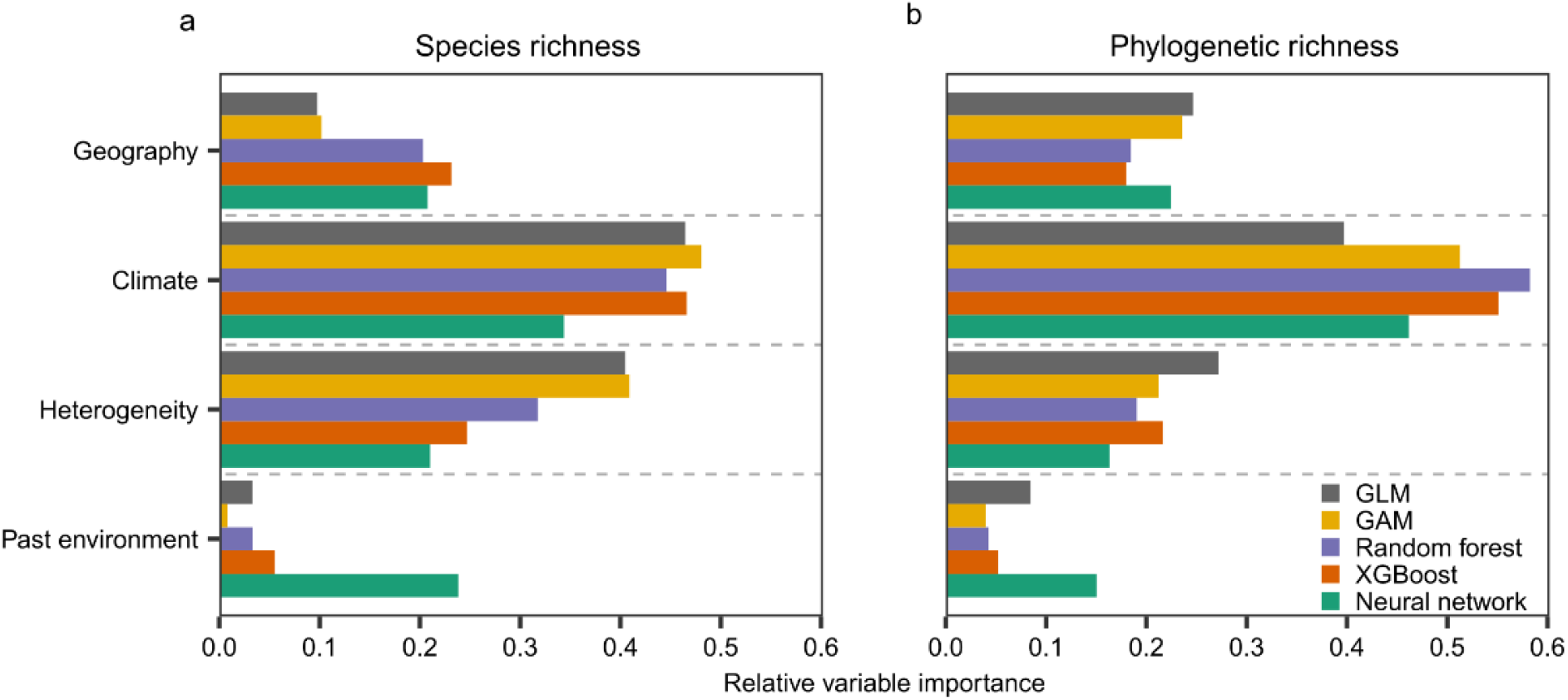
Relative importance of environmental variable categories for explaining global patterns of vascular plant diversity across five non-spatial models. a, species richness; b, phylogenetic richness (Faith’s PD). Relative importance for different variable categories (scaled to sum up to one) was calculated as one minus the Spearman rank correlation coefficient between predictions of the model using a dataset where the values of the predictors of interest were permuted and predictions using the original dataset. Categories of environmental variables are shown in Supplementary Table 1. For the importance of individual environmental variables, see Supplementary Fig. 16.

Geographic variables (area and geographic isolation) explained 9.8-23.1% of the variation in species richness and 18.0-24.6% in phylogenetic richness. Larger regions tend to have higher *in-situ* speciation rates owing to more opportunities for geographic isolation within a region, and lower extinction rates due to larger populations^27,46,47^. These effects should be most pronounced in self-contained, isolated regions like islands, mountains, or other isolated habitats, and less so in regions that are similar to their surroundings^48^. Additionally, larger regions often provide a greater variety of habitats, offering more environmental niches to be occupied by species. Geographic isolation, measured here as the proportion of surrounding landmass, did not explain much variation (0.0-3.9% in species richness; 0.5-3.5% in phylogenetic diversity; Supplementary Fig. 16) for both diversity facets, possibly because our dataset consisted mainly of mainland regions (93.4% of all regions). While geographic isolation is a main driver of insular plant diversity^49^, isolation and peninsular effects seem to play only a minor role on the mainland, where geographic isolation can be expected to be more important for compositional uniqueness of regions and endemism, rather than for richness^18^.

We hypothesized that higher plant diversity would accumulate in regions with long-term climate stability because of low extinction and high speciation rates^16,30^. We therefore assessed the effects of temperature stability and biome variation as proxies for past climatic change for two paleo-time periods, i.e. the last glacial maximum (LGM) and the mid-Pliocene warming period. In contrast to the expected legacy effects of historical variables on modern plant diversity, past environmental conditions only contributed 0.8-5.5% to explaining species richness in most of our models, but up to 23.8% in neural networks. Likewise, past environmental conditions showed higher explanatory power (15.0%) for phylogenetic richness in neural networks than in other models (4.0%-8.5%). Models including spatial trend surfaces or discrete biogeographic regions (i.e. floristic kingdoms) to account for regional idiosyncrasies (after statistically controlling for current and past environments) further improved model fits (Table 1 and Supplementary Table 2). This suggests that in addition to climate stability since the LGM or mid-Pliocene warm period, biogeographic history pre-dating the Pliocene or regional idiosyncrasies other than climatic changes affected modern plant diversity. These historical regional effects are possibly due to dispersal barriers and idiosyncratic colonisation and diversification histories^50,51^.

### Improved global plant diversity maps

We produced global diversity maps for species richness and phylogenetic richness of vascular plants, based on individual well-performing models and model ensembles. Because of its outstanding predictive power and its ability to handle missing data, we consider XGBoost (that includes geographic coordinates) the most powerful single model for predicting plant diversity (Supplementary Figs. 17d and 18d). In addition, we present ensemble predictions which reduce the uncertainty introduced by the choice of one particular modelling technique and therefore improve prediction accuracy^52^. Including region area and its interactions with other predictor variables allowed us to predict plant diversity across global grids of equal area and equidistant hexagons of different grain sizes (i.e. 7,774; 23,322; 69,967 and 209,903 km^2^; Supplementary Figs. 19 and 20). All model predictions and their uncertainties are accessible at https://gift.uni-goettingen.de/shiny/predictions/.

Our ensemble predictions (Fig. 2a, d) describe the global patterns of species and phylogenetic richness with an unprecedented detail and accuracy. The maps capture how diversity varies along environmental gradients and identify global centres of plant diversity (Fig. 2b, e). The highest concentrations of plant species and phylogenetic richness are predicted in Central America, southern Mexico, Andes-Amazonia, the Caribbean, south-eastern Brazil, the Cape region of Southern Africa, Madagascar, Malay Archipelago, Indochina and southern China (Fig. 2b, e), which is in line with empirical observations and previous studies^6,53,54^. While patterns of phylogenetic richness closely resembled species richness (Pearson’s r = 0.97), discrepancies occurred, for example, around the Mediterranean, in Central America, the Caucasus and Himalayas (Supplementary Fig. 21). Differences might result from unequal taxonomic efforts (e.g. many closely related species described separately in Europe) or the uneven distribution of evolutionarily old or young clades across the globe^55,56^. The former suggests that predictions of phylogenetic diversity provide a taxonomically less biased representation of global plant diversity patterns.

**Fig. 2.**
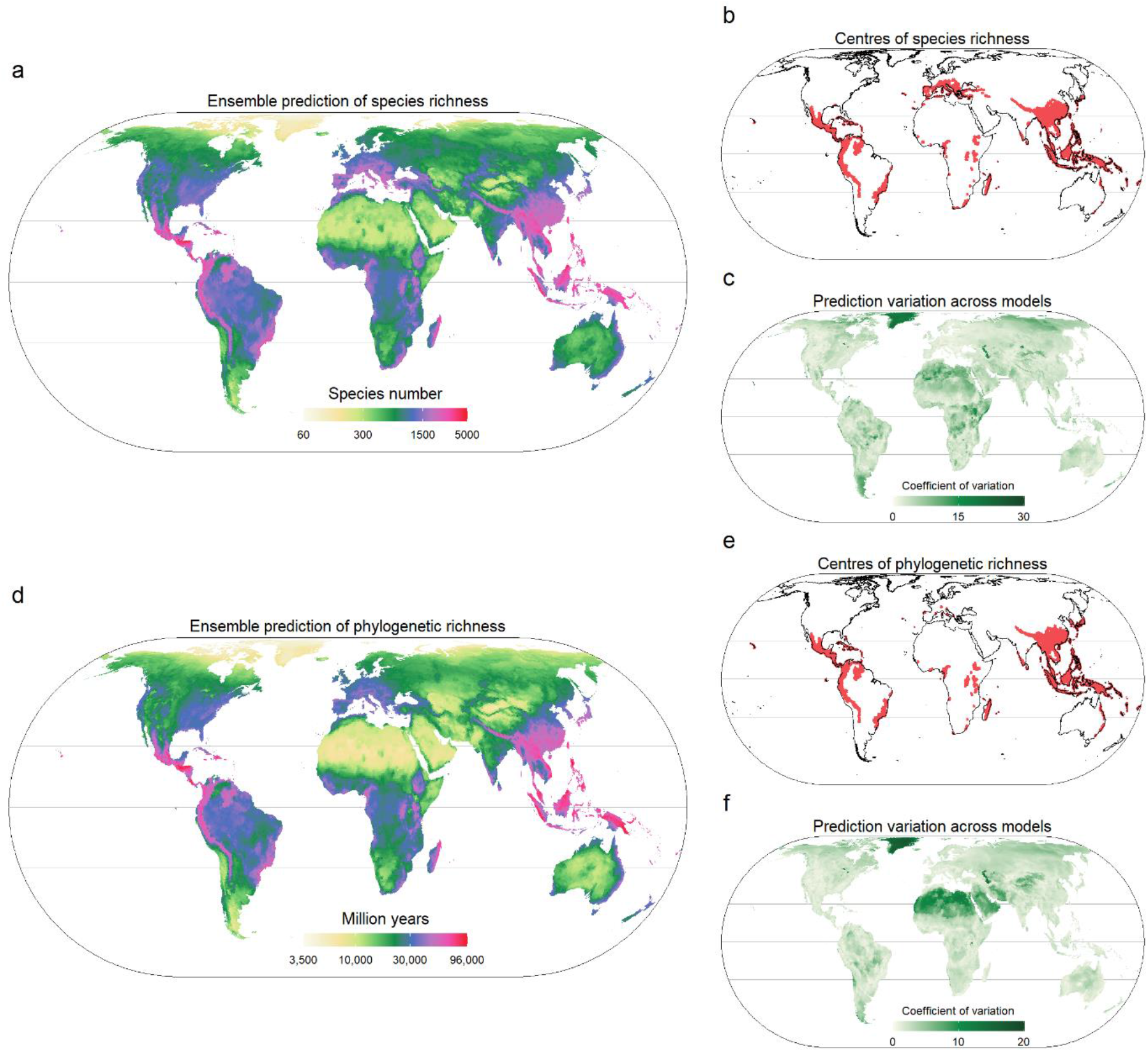
Global patterns of vascular plant diversity predicted across an equal area hexagon grid of 7,774 km^2^ resolution. Species richness (a) and phylogenetic richness (Faith’s PD, d) based on an ensemble of five different models (i.e. three spatial models using machine learning methods, a spatial GAM, and a non-spatial GLM with interactions) weighted by model accuracy; Species richness (b) and phylogenetic richness (e) centres defined as regions with predicted richness values higher than the 90^th^ quantile of the predictions (i.e. containing at least 1,765 plant species and 41,866 Ma of phylogenetic richness per 7,774 km^2^).; Variation of predictions across models used for the ensemble predictions calculated as coefficient of variation of predicted values for species richness (c) and phylogenetic richness (f). Horizontal lines depict the equator and borders of the tropics. In a, b, d, e, log_10_ scale is used and all maps use Eckert IV projection. For maps of all different models and resolutions and data download, see https://gift.uni-goettingen.de/shiny/predictions/.

Thanks to the high-resolution environmental data and modelling techniques that account for complex interactions, regions with steep elevational gradients show finer tuned variation in predicted effects presented here than in previous studies^6,53^. For example, the eastern slopes of the Andes, southern Himalayan slopes, or the northern Kunlun Mountains in China show a finer differentiation from adjacent dryer and less diverse regions than in Kreft and Jetz^6^. At the same time, our ensemble predictions show relatively high values in species-poor regions like non-glaciated parts of Greenland or the Sahara. Here, and in other regions with extreme values of plant diversity, individual models perform better than the ensemble model (Supplementary Figs. 17 and 18), which tends to attenuate extreme values. Besides the important differences just outlined, the ensemble predictions presented here were strongly correlated with model predictions in Kreft and Jetz^6^ (Pearson’s r = 0.872; Supplementary Fig. 22). Apart from the different modelling techniques used and how they account for complex and interactive diversity– environment relationships, differences with previous maps may derive from the accumulation of knowledge on plant diversity worldwide and the continuously updated species distribution data in GIFT used for modelling.

Regions with high species and phylogenetic richness were found to be distributed mostly in mountainous regions (Supplementary Fig. 23). Specifically, tropical mountain ranges, including the tropical Andes, eastern African highlands and various Asian mountains (e.g. in southern China and the Malay Archipelago), are the global centres of plant diversity. The high diversity of tropical mountain ranges is linked to warm and wet climates and heterogeneous environments ^57^. Multiple biogeographical and evolutionary processes, including speciation, dispersal, and persistence that are driven by long-term orogenic and climatic dynamics in mountains have led to outstanding regional plant diversity^57,58^. Orogenic processes constantly change soil composition, nutrient levels and local climate of mountainous regions, thus creating novel and heterogeneous habitats where plant lineages diversify and colonise from neighbouring areas^57^. Moreover, climatic fluctuations stimulate diversification by driving dynamic shifts in habitat connectivity within mountains^58^. Due to their steep environmental gradients and heterogeneous nature, mountain regions provide refugia in times of unfavourable climate^58,59^.

Differences among models (measured as coefficient of variation) were greatest in regions with extreme environments, such as deserts and Arctic regions (Fig. 2c, f). Arctic regions also consistently showed the highest prediction uncertainty across models (Supplementary Figs. 24 and 25). The uncertainties in regions with extreme environments probably stem from two sources. First, extremely species-poor regions might be less well represented in published diversity data. Regions with extreme environments are often part of larger, artificially delimited regions instead of being sampled individually. Those artificially delimited regions are more environmentally heterogeneous, which blurs the signal of extreme environments. Machine learning models are known to not extrapolate well under such conditions^60^. Second, even for regions with relatively homogeneous environments, checklists and Floras do not only include information on predominant but also azonal vegetation, making them appear richer than expected from their prevailing conditions.

## Conclusions

We present the most accurate and comprehensive predictive global maps of vascular plant species richness and phylogenetic richness available to date. They are based on significantly improved global models using comprehensive global inventory-based plant distribution data, high resolution past and current environmental information, and advanced machine learning models. Our findings illustrate that machine learning methods applied to large distribution and environmental datasets help to disentangle underlying complex and interacting associations between the environment and plant diversity. Machine learning methods therefore help to improve both the fundamental understanding and quantitative knowledge in biogeography and macroecology. The updated global diversity maps of vascular plant diversity at multiple grain sizes (available at https://gift.uni-goettingen.de/shiny/predictions/) provide a solid foundation for large-scale biodiversity monitoring and research on the origin of plant diversity, and subsequently support future global biodiversity assessments and environmental policies.

## Methods

### Species distribution data and species richness

To calculate species richness and phylogenetic richness, we used the species composition of native vascular plants in regional checklists and Floras from the Global Inventory of Floras and Traits^10^ (GIFT version 2.1: http://gift.uni-goettingen.de). In GIFT, all non-hybrid species names are standardized and validated based on taxonomic information provided by The Plant List (version 1.1, http://www.theplantlist.org) and additional resources available via iPlant’s Taxonomic Name Resolution Service(TNRS)^10,61^. The original database contains > 3000 geographic regions representing islands, protected areas, biogeographical regions and political units (e.g. countries, provinces in China or states in the USA). We excluded regions with incomplete native vascular plant checklists, incomplete data for predictor variables, or an area of less than 100 km^2^. Furthermore, we coped with overlapping regions in two steps. First, for those overlapping regions from one individual literature source, we only used non-overlapping regions preferring smaller over larger regions (e.g. the individual states of Brazil instead of the entire country). Second, for overlapping regions from different literature sources, we removed the larger regions. If smaller regions covered only parts of the larger regions, we retained both smaller and larger regions. A total of 298,087 vascular plant species from 830 geographic regions, consisting of 775 mainland regions and 55 islands or island groups was used to calculate species richness (i.e. taxonomic richness). The geographic regions in the dataset were distributed representatively across the entire globe, covering all major biomes (Supplementary Fig. 1).

### Phylogeny reconstruction and phylogenetic richness

We used a large, dated megatree of vascular plants, GBOTB_extended^42^, as a backbone to generate a phylogeny for all species in the dataset. The megatree was derived from the GBOTB tree for seed plants by Smith and Brown^62^ and the phylogeny for pteridophytes in Zanne et al.^63^. Because of missing phylogenetic information for undescribed species, we excluded them from the dataset for calculating phylogenetic richness, leading to a dataset including 295,417 species in 466 families of vascular plants. All families and 10,128 out of 14,962 genera (67.7%) in the dataset were included in the megatree. We bound the remaining genera and species into their respective families and genera using the “Scenario 3” approach in the R package *V.PhyloMaker*. In “Scenario 3”, the weighted positioning of the additional taxa depends on the length and amount of already existing tips per taxon^42^. 91.95% out of the 295,417 species in the dataset were from genera already present in the backbone.

Several indices exist for capturing different dimensions of phylogenetic diversity including richness, divergence and regularity^64^. Here, we focus on phylogenetic richness, which represents the amount of unique phylogenetic history present in an assemblage^64^. We chose Faith’s PD, a common measure of phylogenetic richness, calculated as the sum of the branch lengths of all species coexisting in a region^41^, which is directly comparable to species richness. Even though highly correlated to species richness, we refrained from standardising phylogenetic richness as we were not interested in the deviation of phylogenetic richness from expectations based on species richness, but rather as a richness metric accounting for phylogenetic relationships that is comparable to species richness.

A recent study^65^ showed that phylogenetic richness derived from a phylogeny resolved only at the genus level was nearly perfectly correlated with phylogenetic richness derived from a phylogeny resolved fully at the species level (Pearson’s r: 0.997-1). This suggests that patterns of phylogenetic richness would be similar regardless of whether the phylogeny used to calculate phylogenetic richness is resolved at the genus or species level. As a sensitivity analysis, we disregarded species from genera that were absent from the phylogenetic backbone and constructed a tree only for the remaining species to test for the effect of missing genera (i.e. additional genera bound to their respective families) on phylogenetic richness. We found that phylogenetic richness calculated from the tree resolved at the genus level was nearly perfectly correlated to phylogenetic richness based on the tree with missing genera added (Pearson’s r = 0.998). This suggests that patterns of phylogenetic richness would be equal regardless of whether the phylogeny used is fully resolved at the genus level or still included unresolved genera. In the following analyses, we used phylogenetic richness derived from the tree, including all 295,417 species.

### Predictor variables

We identified a set of candidate predictor variables hypothesised to affect plant distributions and diversity and classified them into four categories: geography, climate, environmental heterogeneity and past environmental conditions. Twenty-two predictors were considered in the original dataset (Supplementary Table 1). These have been shown or hypothesised to contribute to geographic patterns of plant diversity in previous studies^6,20,32,33^. Geographic variables were region area (km^2^) and the summed proportion of landmass area in the surrounding area of the target region within buffer distances of 100 km, 1000 km, 10,000 km, serving as a measure of geographic isolation^49^. Climatic variables included 13 biologically relevant temperature and precipitation variables. These variables represent annual averages, seasonality and limiting climatic factors (e.g. length of growing season, precipitation of warmest quarter), capturing the main aspects of climate important for plant diversity^66,67^. Furthermore, gross primary productivity^68^ was included as a measure of potential plant productivity based on available solar energy and water. Climatic variables were extracted as mean values across the input raster layers per region. The number of soil types^36^ and elevational range^69^ were calculated for each region as proxies for environmental heterogeneity within regions.

To determine the contribution of past environmental conditions to modern diversity patterns, we calculated biome area variation since the Pliocene and the Middle Miocene, temperature anomaly since the mid-Pliocene warm period, temperature stability since the last glacial maximum (LGM) and velocity of temperature change since the LGM. Terrestrial biomes are affected by multiple drivers containing atmospheric circulation, precipitation and temperature patterns, and thus changes in biome distributions represent major environmental changes through geological time. To calculate biome area variation, we used biome distribution maps at present^70^, the LGM (∼25 – 15 ka)^71^, the mid-Pliocene warm period (mid-Piacenzian, ∼3.264 – 3.025 Ma)^72^ and the Middle Miocene (∼17–15 Ma)^73^. The LGM represents a cooler period compared to present-day conditions, characterized by large glaciated areas, expanded dry deserts and reduced forest biome areas. On the contrary, the mid-Pliocene and the Middle Miocene are two relatively warm periods compared to the present-day, characterised by a decreased ice loading of the continents and expanded forest biomes at the expense of deserts^72,73^. Biome definitions differed across the four datasets. We therefore regrouped biomes to match across datasets (Supplementary Table 3). We extracted each biome’s area inside each region at each time slice and standardised it by dividing it by the area of the region. Then we calculated Euclidean distances of biome area change between every two adjacent time-slices for each region and averaged the distance across all time-slice periods. We acknowledge potential drawbacks of this approach due to the coarse resolution and uncertainty of the original past biome maps. Because of the coarse resolution of the Middle Miocene map and absent data for some geographic regions in this map, we only used biome area variation since the Pliocene and excluded Miocene biome variation from further analyses.

In addition, we calculated temperature stability from two paleo-time periods until present, i.e. the LGM (∼ 21 ka) and the mid-Pliocene warm period, representing cooler and warmer climates compared to the current climate, respectively. Temperature stability since the LGM was calculated using the *climateStablity* R package. It calculates temperature differences between 1000 year time slices expressed as standard deviation and averages the results across all time slices. The stability is then the inverse of the mean standard deviation rescaled to [0,1]^74^.

Temperature anomaly since the mid-Pliocene was calculated as the difference in annual mean temperature between the mid-Pliocene warm period and present-day^75^. The velocity of temperature change since the LGM was calculated as the ratio between temporal change and contemporary spatial change in temperature, representing the speed with which a species would have to move its range to track analogous climatic conditions^76^.

An alternative way to evaluate effects of biogeographic history on plant diversity is to account for predefined discrete geographic regions influencing diversity via differences in diversification history and dispersal barriers. We therefore included floristic kingdoms^77^ as an additional categorical variable in the models and compared the performance of models with and without floristic kingdom to assess if we managed to model the effect of biogeographic history sufficiently by only including the variables that directly quantify past environmental change.

### Statistical models

To quantify diversity–environment relationships, we fitted five different types of models with species richness and phylogenetic richness as response variables: generalised linear models (GLMs), generalised additive models (GAMs), random forests, extreme gradient boosting (XGBoost) and neural networks. To compare model performance across model types, we used the same set of predictors across models. Since there was significant collinearity between the 22 predictors in the initial dataset, we removed variables with low contribution to predictions until the variance inflation factors (VIF) of all remaining variables was below a threshold of five. The contribution to predictions was based on a preliminary ranking of predictor variables using random forests and a stepwise forward strategy for variable introduction^78^. Like this, we selected a subset of 15 predictor variables minimising redundancy and maximizing model performance to fit models (bold in Supplementary Table 1; Supplementary Fig. 26). The predictors retained represented all aspects (geography, climate, environmental heterogeneity and past environment) that could potentially affect plant diversity patterns.

To perform GLMs and GAMs, we used a negative binomial error distribution with a log-link function for species richness to cope with over-dispersion of the response variables, and a Gaussian error distribution with log-link function for phylogenetic richness. For the GLMs, some predictors were log-transformed owing to their skewed distribution (i.e. area, temperature seasonality, number of wet days, precipitation seasonality, precipitation of warmest quarter, gross primary productivity, elevational range, number of soil types and velocity in temperature since LGM). All continuous predictor variables were standardised to zero mean and unit variance to aid model fitting and making their parameter estimates comparable. Although fitting GLMs with 15 predictors might seem excessive, it is suggested not to exclude predictors hypothesized to be important when collinearity is minimised and not a hindrance to analysis^79^. Thus, in our GLMs, we built the full model including 15 predictors and then simplified the model using Akaike’s information criterion (AIC). Predictors were tested in turn, and removed if AIC values were larger in the complex models compared to the simpler ones (Supplementary Table 2). In GAMs, we used penalised regression smoothers (with 9 spline bases for species richness and 10 spline bases for phylogenetic richness) for each predictor to estimate the smooth terms. The number of spline bases were selected from values between 2 and 10 using cross validation to optimise model performance (i.e. minimising root mean square error). Additionally, we used a gamma value of 1.4 to reduce overfitting without compromising model fit^80^ and also included a double penalty to variable coefficients. We used the R packages *MASS* to fit negative binomial generalised linear models and *mgcv*^80^ to fit GAMs.

In addition, we applied machine learning techniques, i.e. random forests, extreme gradient boosting and neural networks, to fit global models of plant diversity. For these machine learning methods, species richness and phylogenetic richness were log-transformed prior to modelling to reduce skewness of their distributions. A set of tuning parameters (i.e. hyperparameters), which cannot directly be estimated from the data, needs to be set beforehand. These hyperparameters determine the training strategy and related efficiency of the algorithms. It is commonly suggested to tune hyperparameters to maximise model performance before running models for a certain problem^81^. We used the *train* function from the R package *caret* to optimise the model tuning parameters for the three machine learning models used here^82^. We used repeated cross-validation and selected the parameters that produced the lowest root mean squared error. We then refitted the final models using these optimal parameters. The R packages *ranger* was used to fit random forests^83^, *xgboost* to fit XGBoost^84^ and *neuralnet* to fit neural networks^85^.

Random forests are an ensemble learning method that builds a large collection of decision trees and outputs average predictions of the individual regression trees. After tuning the hyperparameters, we used 8 variables randomly sampled as candidates at each split and each node contained at least 4 samples (minimal node size) for species richness, while 6 variables were randomly sampled at each split and each node contained at least 4 samples for phylogenetic richness; we fitted 500 regression trees. Extreme Gradient Boosting (XGBoost) is a scalable end-to-end machine learning system for tree boosting. Like other tree boosting methods, XGBoost is an ensemble model of decision trees trained sequentially; it learns decision trees by fitting the residual errors in each iteration. Several innovations make XGBoost highly effective, including a novel tree learning algorithm for handling sparse data, and a theoretically justified weighted quantile sketch procedure enabling handling instance weights in approximate tree learning^84^.

There are three types of parameters that have to be set in XGBoost: general parameters, task parameters and booster parameters. General parameters are related to which booster is used, and here we selected tree-based models. For task parameters, we chose regression with squared loss for ranking tasks. Booster parameters define how to build the tree models. In our resulting XGBoost model for species richness, we used the booster parameters max_depth = 4, eta = 0.1, gamma = 0, colsample_bytree = 0.8, min_child_weight = 1 and subsample = 0.7. In the XGBoost model for phylogenetic richness, we used the booster parameters max_depth = 4, eta = 0.1, gamma = 0, colsample_bytree = 0.7, min_child_weight = 1 and subsample = 0.8. Further, we set the maximum number of boosting iterations to 200. Both random forests and XGBoost control for overfitting by utilising ensemble strategies, i.e. bagging and gradient boosting. While column (feature) sampling is used by both methods to further prevent overfitting, XGBoost utilises two additional techniques, regularisation and shrinkage^84^.

Neural networks are a machine learning method that comprises a collection of connected units (neurons) and their connections (edges). We applied feed-forward neural networks with three hidden layers between the input and output layer. We used resilient backpropagation with the weight backtracking algorithm to compute the neural networks. Compared with traditional backpropagation algorithms, this algorithm applies a separate learning rate, which can be changed during the training process^85^. Moreover, we applied logistic activation. We scaled the response after log-transformation and all predictors to fall within a range of 0 to 1 to improve neural network stability and modelling performance. For every hidden layer, we trained the number of neurons, and the architecture of the final networks for prediction was (8,4,8) in the species richness model and (8,4,10) in the phylogenetic richness model.

Species distribution data and environmental predictors are often spatially autocorrelated. On the one hand, this may lead to biased parameter estimates which needs to be accounted for, and, on the other hand, including spatial information in models may increase their predictive power^14,43^. Because of this, we generated spatial models using different modelling techniques. To account for spatial autocorrelation in GLM residuals, we used simultaneous autoregressive (SAR) models, a common method used to deal with spatial autocorrelation in ecological data. Simultaneous autoregressive models can take three different types depending on where the spatial autoregressive process is believed to occur. Here we selected models of the spatial error type, which is recommended for use when dealing with spatially autocorrelated species distribution data^6,86^. We evaluated SAR models with different neighbourhood structures and spatial weights (lag distances between 200 and 3000 km, weighted and binary coding). As the final simultaneous autoregressive model, we chose a model with weighted neighbourhood structure and 800 km lag distance for both species richness and phylogenetic richness, which had the minimal AIC and the best reduction of spatial autocorrelation in the residuals. Species richness and phylogenetic richness were log-transformed prior to modelling. In GAMs, we added a two-dimensional spline on geographical coordinates, which accounts for spatial autocorrelation in model residuals^14,43^. In machine learning, to cope with spatial autocorrelation, we fitted spatial models respectively for the three machine learning techniques by including cubic polynomial trend surfaces (i.e. latitude (Y), centred longitude (X) as well as X^2^, XY, Y^2^, X^3^, X^2^Y, XY^2^ and Y^3^)^87,88^. Overall, the spatial models successfully removed spatial autocorrelation from model residuals (Supplementary Fig. 27).

To compare our models to published global models of plant species richness, we rebuilt these models for the data set analysed here. First, we fitted the best model from Kreft and Jetz^6^, a combined six-predictor model using GLM; and second, we built a GAM using the same model structure as Keil and Chase’s smooth model^14^, which contained a two-dimensional spline on geographical coordinates, 15 single predictors and interactions between each individual predictor and area. We ran models including the same 15 predictor variables and floristic kingdom using random forests and XGBoost, and compared them with the models without floristic kingdom.

Adding floristic kingdom increased collinearity between predictors. However, the two tree-based models are able to handle multicollinearity when they are used for prediction. Random forests in the *ranger* R package can handle categorical variables automatically; however, XGBoost only works with numeric vectors. We therefore converted all other forms of data into numeric vectors. Here we used one-hot encoding (0,1) to convert the floristic kingdom into dummy variables for the XGBoost model.

To estimate the relative importance of each environmental predictor, we used a consistent method across model types. We randomly reshuffled values of the predictor of interest in the dataset, predicted the response variables based on the modified dataset and calculated the Spearman rank correlation coefficient between those predictions and the predictions using the original dataset. The relative importance of the predictor of interest was calculated as one minus the correlation coefficient divided by the sum of one minus the correlation coefficients of all predictors^89^. Likewise, to compare the relative importance of different categories of predictor variables (categories in Supplementary Table 1), we permuted values of a subset of predictors belonging to one category, correlated the predictions of the model using the modified dataset and predictions using the original dataset, and estimated the importance of each category as one minus the Spearman rank correlation coefficient divided by the sum of one minus the correlation coefficients of all predictor categories.

We used partial dependence plots from the R package *pdp* to visualise the relationships between diversity metrics and the single predictors across models. Partial dependence plots visualise the relationship between a subset of the predictors (typically 1-3) and the response while accounting for the average effect of all other predictors in the model^90^. To identify and visualise important two-way interactions between predictor variables, we followed Lucas^91^. We first calculated the interaction importance for each covariate by decomposition of the prediction models^92^. Then we calculated the two-way interaction strengths between the covariates of interest and all other covariates. Finally, we visualised important two-way interactions using two-predictors partial dependence plots. Contrary to the machine learning methods, interactions in GLMs need to be defined prior to modelling. We fitted GLMs including energy-water, energy-environmental heterogeneity and area-environment interactions, as found to have strong effects on diversity patterns in our machine learning models or suggested by previous studies^6,14,20,33^. The models were then simplified based on AIC values (Supplementary Table 2).

### Cross-validation

To assess the accuracy of model predictions across all different model types, we used random 10-fold cross-validation and spatial 68-fold cross validation. In random cross validation, the observations in the dataset were randomly partitioned into 10 nearly equally-sized sets. Nine sets were used to train the model, which was then used to make predictions for the remaining set. The predictions were then compared to observed values. This process was repeated until all 10 sets had been predicted. For spatial cross validations, observations were grouped into spatially homogeneous clusters if their pairwise geographic distances were smaller than the threshold of spatial autocorrelation to remove potentially spatial dependence between training and test data^93^. Spatial clusters were generated using a hierarchical cluster analysis (complete linkage method) of the distance matrix of observed geographical coordinates and a clustering height (i.e. the threshold of spatial autocorrelation). The threshold of spatial autocorrelation (i.e. the maximum distance between regions within each cluster) was defined here as 2,000 km based on the spatial correlograms of raw observed species and phylogenetic richness, where the Moran’s I almost reached zero (Supplementary Fig. 27).

To quantify model predictive performance, we summarised the cross validation results using root mean squared error and two different pseudo-coefficients of determination to quantify the amount of variation explained by the model based on out-of-bag samples. R^2^_CORR is the coefficient of determination of a linear model of the predicted and observed values from all repetitions of the cross-validation. R^2^_Accuracy is the amount of variation explained by the model, calculated as R^2^_Accuracy =[1-SSE/SST]^36^, where SSE is the sum of the squared error between observation and prediction and SST is the total sum of squares. The model with the lowest root mean squared error and highest R^2^_CORR/ R^2^_Accuracy was identified as the best predictive model. For all models, we calculated cross-validation results for log-transformed observed and predicted species and phylogenetic richness, because species richness and phylogenetic richness were log-transformed prior to modelling for machine learning models and fitted with log link functions in GLMs and GAMs.

In our models, variation explained based on spatial cross-validation was consistently lower than variation explained based on random cross-validation, likely because the former excludes subsets of regions with specific environmental characteristics and biogeographic histories from the training data and is therefore less representative of the globe and its environmental spectrum.

### Predictions

We used the resulting models to predict vascular plant species and phylogenetic richness across global grids of four different resolutions (i.e. 7,774; 23,322; 69,967 and 209,903 km^2^ hexagon size). We used the *dggridR* R package to produce a grid of equal-area and equidistant hexagons across the Earth’s surface clipped for global coastlines. Islands smaller than 1.5 times the gridcell size were treated as entire entities instead of subdividing them into several partial grid cells. For each hexagon, we calculated the same predictor variables as for the geographic regions used for fitting the models. We then used the models to predict vascular plant species richness and phylogenetic richness, and mapped the predictions across the hexagon grid. Seven predictors (i.e. surrounding landmass proportion, gross primary productivity, mean temperature of growing season, biome area variation since Pliocene, temperature stability since LGM and temperature anomaly since the mid-Pliocene warm period in all resolutions; precipitation seasonality in 7,774 and 23,322 km^2^ resolution) had missing values in a small number of hexagons at the margins of continents, northern Africa and Greenland. XGBoost can handle missing data when it is used for predicting. An optimal default direction in each tree node is learned from the trained data in the model constructing process. When there are missing values in predictor data, the observation is classified into the default branch. Random forests, neural networks, GLMs and GAMs cannot deal with missing values. We therefore interpolated predictor values for hexagons from their neighbours. To interpolate biome area change, we defined the biome type of each hexagon where missing values occurred in the Pliocene map according to their neighbouring cells. Besides predictions based on individual models, we used an ensemble prediction procedure, which averages the predictions based on the models fitted by different techniques weighted by model accuracy (the inverse of the model squared error) from the random cross-validation process^52^. As centres of plant diversity based on the ensemble predictions, we defined regions with predicted richness values higher than the 90^th^ quantile, i.e. containing at least 1,765 plant species and 41,866 Ma of phylogenetic richness at a resolution of 7,774 km^2^.

### Uncertainty

To assess variation of the predictions across models, we calculated the coefficient of variation of predicted values for each hexagon grid cell. The coefficient of variation is defined as the ratio of the standard deviation to the mean, which accounts for the differences in diversity between regions and thereby avoids artificially high uncertainty of high diversity regions. Additionally, we calculated standard errors of predictions for GLMs, GAMs and random forests. For XGBoost and neural networks, we modelled the relationship between model residuals and environmental predictors from the raw data, and used this model to predict uncertainty across the hexagon grids.

## Data availability

Predictions of vascular plant species and phylogenetic richness and model uncertainties based on the various statistical models applied here are available at https://gift.uni-goettingen.de/shiny/predictions/.

## Acknowledgements

L.C. was supported by China Scholarship Council (CSC) Grant (No.201808330443). P.P. was supported by EXPRO grant no. 19-28807X (Czech Science Foundation) and long-term research development project RVO 67985939 (Czech Academy of Sciences). M.W. acknowledges DFG funding via iDiv (DFG FZT 118, 202548816). We thank Alexandr Ebel and Christian König for contributing data and discussions about the manuscript.

## Author contributions

L.C., H.K., and P.W. conceived of the idea and developed the conceptual framework of the study. All authors were involved in collecting the data. L.C. performed the statistical analyses and wrote the first draft of the manuscript with the help of H.K., A.T. and P.W. All authors contributed to the writing and interpretation of the results.

