## Supplementary material for "Machine learning improves global models of plant diversity"

### Supplementary material for Cai et al. (2020), Machine learning improves global models of plant diversity, bioRxiv.

**Supplementary Table 1 | List of environmental predictor variables hypothesized to affect plant diversity patterns.** Abbreviations in bold highlight predictor variables that were retained after the initial predictor-selection process to limit collinearity and used to fit different models of species richness and phylogenetic richness of vascular plants. References can be found in Supplementary References 1.

| Category | Abbreviation | Variable description | Unit | Resolution | Reference |
|---|---|---|---|---|---|
| Geography | **Area** | Area of geographic regions | $km^2$ | - | 1 |
| Geography | **SLMP** | Surrounding landmass proportion: Summed proportions of landmass area surrounding the target region within buffer distances of 100 km, 1000 km and 10,000 km | - | - | 2 |
| Climate | AMT | Annual mean temperature | °C | 30 arc-seconds | 3 |
| Climate | PET | Potential evapotranspiration | mm | 30 arc-seconds | 4 |
| Climate | **LengthGrow** | Length of growing season (days): Number of days with temperatures exceeding a threshold of 0.9°C, without snow cover, and with sufficient soil water | n | 30 arc-seconds | 5 |
| Climate | **MeanTempGrow** | Mean temperature of growing season | °C | 30 arc-seconds | 5 |
| Climate | MeanT_WetQ | Mean temperature of wettest quarter | °C | 30 arc-seconds | 3 |
| Climate | IS | Isothermality, quantifying how large the diurnal variation in temperature is proportional to the annual variation: The ratio of the mean diurnal range divided by the annual temperature range, and then multiplied by 100. | - | 30 arc-seconds | 3 |
| Climate | **TS** | Temperature seasonality: standard deviation of mean monthly temperature *100 | °C | 30 arc-seconds | 3 |
| Climate | TAR | Temperature annual range | °C | 30 arc-seconds | 3 |
| Climate | AP | Annual precipitation | mm | 30 arc-seconds | 3 |
| Climate | **Wetdays** | Number of days per year with precipitation >0.1mm | n | 10 arc - minute | 6 |
| Climate | **PrecipWarmQuater** | Precipitation of warmest quarter | mm | 30 arc-seconds | 3 |
| Climate | PrecipGrow | Precipitation of growing season | mm | 30 arc-seconds | 5 |
| Climate | **PS** | Precipitation seasonality: Ratio of the standard deviation of the monthly total precipitation to the mean monthly total precipitation (coefficient of variation in monthly precipitation) | - | 30 arc-seconds | 3 |
| climate | **GPP** | Gross primary productivity | gcarbon/$m^2$ | 30 arc-seconds | 7 |
| Heterogeneity | **Elev_range** | Elevational range: absolute difference between highest and lowest elevation within a given area | m | 30 arc-seconds | 8 |
| Heterogeneity | **Soildiv** | Number of different soil types based on World Reference Base classification system | n | 30 arc-seconds | 9 |
| Heterogeneity | Homogeneity | Homogeneity: similarity of MODIS enhanced vegetation index between adjacent pixels | - | 30 arc-seconds | 10 |
| Past environments | **VelocityTemp_LGM** | Climate change velocity in temperature for the last 21,000 years (since the Last Glacial Maximum) calculated as the ratio between temporal change and contemporary spatial change in temperature | m/yr | 30 arc-seconds | 2,11 |
| Past environments | **TempStability_LGM** | Temperature stability for the last 21,000 years (since the Last Glacial Maximum): Mean standard deviation in annual mean temperature between time slices of 1000 years each over the last 21,000 years; "stability" is defined as the inverse of this deviation re-scaled here between 0 and 1. | - | 2.5 degrees | 12 |
| Past environments | **TempAnomaly_midPliocene** | Difference for annual mean temperature between the mid-Pliocene warm period (~3.264-3.025 Ma) and the present-day, i.e. the value of past minus present. | °C | 2.5 arc-minutes | 13,14 |
| Past environments | Biome_Miocene | Euclidean distance of biome area changes across four time periods (present, Last Glacial Maximum, Pliocene and Middle Miocene) | - | - | 15–18 |
| Past environments | **Biome_Pliocene** | Euclidean distance of biome area changes across three time periods (present, Last Glacial Maximum and Pliocene) | - | - | 15,17,18 |
| Past environments | Kingdom | Floristic kingdoms: Antarctic kingdom, Australis kingdom, Cape kingdom, Holarctic kingdom, Neotropic kingdom, Paleotropic kingdom. | factor | - | 19 |

**Supplementary Table 2 | Model assessment results.** Each model was assessed for its fit based on training data, and for its predictive performance based on all out-of-bag samples using both random 10-fold cross-validation and spatial 68-fold cross-validation. Accuracy statistics are provided on log-scale. Values shown are: root mean squared error (RMSE); the coefficient of determination of a linear model of predicted vs. observed richness (R2_CORR); the amount of variation explained by the model calculated as one minus the ratio of the sum of the squared error between observation and prediction to the total sum of squares (R2_Accuracy).

Non-spatial models were fitted with 15 predictors except Minimum adequate GLM, Full GLM with interaction terms and GLM simplified with interaction terms, while spatial models also contained spatial effects (i.e. GAM included the spline of geographic coordinates and machine learning methods included cubic polynomial trend surfaces). Minimum adequate GLM for species richness contained 9 predictors (i.e. Area, LengthGrow, MeanTempGrow, GPP, TS, PS, Elev_range, Soildiv, Biome_Pliocene; See Table 1 for abbreviations); Minimum adequate GLM for phylogenetic richness contained 13 predictors (PS and wetdays removed). Full GLM with interaction terms was fitted with all 15 environmental predictors and interactions between area and each individual predictor, and between energy and environmental heterogeneity (i.e. MeanTempGrow:Soildiv; MeanTempGrow:Elev_range), as well as energy and water (i.e. MeanTempGrow:PrecipWarmQuater; MeanTempGrow:Wetdays). Simplified GLM with interaction terms for species richness contained 13 individual predictors (PS and TempStability_LGM removed) and 9 interaction terms between area and other predictors and between energy and environmental heterogeneity; simplified GLM with interaction terms for phylogenetic richness contained 14 main predictors (PS removed) and 13 interaction terms between area and other predictors, between energy and environmental heterogeneity, as well as energy and water. GLM (Kreft and Jetz) was a combined six-predictor model as in Kreft and Jetz[20]. SMOOTH Model (Keil & Chase) was fitted using the same model structure as Keil and Chase's smooth model[21], which contained a two-dimensional spline on geographical coordination, 15 individual predictors and the interactions between each individual predictor and area. Likewise, REALM Model was fitted using the same model structure as Keil and Chase's realm model[21], which contained floristic kingdom, 15 individual predictors and the interactions between each individual predictor and area. XGBoost with kingdom and random forests with kingdom were fitted with the same 15 predictors and floristic kingdom.

| Models | Species richness | | | | | | | | | Phylogenetic richness | | | | | | | | |
|---|---|---|---|---|---|---|---|---|---|---|---|---|---|---|---|---|---|---|
| | Model fits | | | Random cross-validation | | | Spatial cross-validation | | | Model fits | | | Random cross-validation | | | Spatial cross-validation | | |
| | RMSE | $R^2$_CORR | $R^2$_Accuracy | RMSE | $R^2$_CORR | $R^2$_Accuracy | RMSE | $R^2$_CORR | $R^2$_Accuracy | RMSE | $R^2$_CORR | $R^2$_Accuracy | RMSE | $R^2$_CORR | $R^2$_Accuracy | RMSE | $R^2$_CORR | $R^2$_Accuracy |
| **Non-spatial models** | | | | | | | | | | | | | | | | | | |
| Full GLM | 0.510 | 0.657 | 0.657 | 0.525 | 0.636 | 0.636 | 0.582 | 0.568 | 0.561 | 0.506 | 0.639 | 0.471 | 0.514 | 0.630 | 0.452 | 0.552 | 0.594 | 0.359 |
| Minimum adequate GLM | 0.511 | 0.657 | 0.657 | 0.520 | 0.643 | 0.643 | 0.548 | 0.609 | 0.608 | 0.506 | 0.640 | 0.470 | 0.513 | 0.631 | 0.454 | 0.548 | 0.600 | 0.369 |
| Full GLM with interaction | 0.457 | 0.722 | 0.722 | 0.478 | 0.695 | 0.694 | 0.535 | 0.627 | 0.620 | 0.397 | 0.697 | 0.661 | 0.419 | 0.668 | 0.623 | 0.467 | 0.616 | 0.531 |
| Simplified GLM with interaction | 0.457 | 0.722 | 0.722 | 0.471 | 0.704 | 0.704 | 0.502 | 0.665 | 0.664 | 0.396 | 0.697 | 0.663 | 0.412 | 0.676 | 0.635 | 0.453 | 0.630 | 0.559 |
| GAM | 0.410 | 0.775 | 0.775 | 0.437 | 0.743 | 0.742 | 0.507 | 0.664 | 0.658 | 0.328 | 0.778 | 0.769 | 0.359 | 0.735 | 0.723 | 0.430 | 0.647 | 0.604 |
| Random forests | 0.403 | 0.786 | 0.775 | 0.415 | 0.774 | 0.761 | 0.511 | 0.642 | 0.639 | 0.309 | 0.807 | 0.795 | 0.317 | 0.796 | 0.784 | 0.395 | 0.669 | 0.667 |
| XGBoost | 0.107 | 0.985 | 0.984 | 0.389 | 0.791 | 0.791 | 0.487 | 0.673 | 0.673 | 0.084 | 0.986 | 0.985 | 0.295 | 0.813 | 0.813 | 0.384 | 0.687 | 0.685 |
| Neural networks | 0.285 | 0.887 | 0.887 | 0.451 | 0.725 | 0.718 | 0.604 | 0.552 | 0.496 | 0.244 | 0.872 | 0.872 | 0.328 | 0.774 | 0.769 | 0.419 | 0.650 | 0.628 |
| **Spatial models** | | | | | | | | | | | | | | | | | | |
| SAR | 0.385 | 0.795 | 0.794 | 0.537 | 0.601 | 0.600 | 0.548 | 0.586 | 0.584 | 0.295 | 0.813 | 0.813 | 0.416 | 0.631 | 0.629 | 0.426 | 0.614 | 0.611 |
| GAM | 0.383 | 0.802 | 0.802 | 0.413 | 0.769 | 0.769 | 0.499 | 0.672 | 0.667 | 0.312 | 0.794 | 0.791 | 0.340 | 0.757 | 0.751 | 0.416 | 0.663 | 0.633 |
| Random forests | 0.384 | 0.810 | 0.795 | 0.398 | 0.796 | 0.780 | 0.502 | 0.660 | 0.653 | 0.292 | 0.828 | 0..816 | 0.303 | 0.815 | 0.803 | 0.379 | 0.697 | 0.694 |
| XGBoost | 0.096 | 0.988 | 0.987 | 0.371 | 0.809 | 0.809 | 0.463 | 0.703 | 0.703 | 0.081 | 0.986 | 0.986 | 0.279 | 0.833 | 0.833 | 0.351 | 0.737 | 0.737 |
| Neural networks | 0.253 | 0.911 | 0.911 | 0.422 | 0.756 | 0.753 | 0.587 | 0.590 | 0.522 | 0.203 | 0.911 | 0.911 | 0.314 | 0.792 | 0.789 | 0.433 | 0.645 | 0.597 |
| **Other models** | | | | | | | | | | | | | | | | | | |
| GLM (Kreft and Jezt 2007) | 0.452 | 0.716 | 0.716 | 0.457 | 0.708 | 0.708 | 0.494 | 0.663 | 0.659 | | | | | | | | | |
| SMOOTH Model (Keil & Chase) | 0.394 | 0.791 | 0.791 | 0.421 | 0.761 | 0.761 | 0.487 | 0.686 | 0.683 | | | | | | | | | |
| REALM Model (Keil & Chase) | 0.415 | 0.770 | 0.770 | 0.445 | 0.734 | 0.733 | 0.494 | 0.679 | 0.673 | | | | | | | | | |
| XGBoost with kingdom | 0.107 | 0.985 | 0.984 | 0.388 | 0.791 | 0.791 | 0.472 | 0.693 | 0.692 | 0.138 | 0.960 | 0.959 | 0.290 | 0.820 | 0.820 | 0.364 | 0.720 | 0.718 |
| Random forests with kingdom | 0.398 | 0.792 | 0.780 | 0.414 | 0.775 | 0.763 | 0.527 | 0.632 | 0.626 | 0.305 | 0.809 | 0.800 | 0.316 | 0.795 | 0.785 | 0.407 | 0.654 | 0.654 |

**Supplementary Table 3 | Homogenization of biome classifications for current maps, Last Glacial Maximum (LGM), Pliocene (mid-Piacenzian) and Middle Miocene.**

| New biome type | Current | LGM | Pliocene (mid-Piacenzian) | Middle Miocene |
|---|---|---|---|---|
| **Tropical forest** | Tropical & subtropical moist broadleaf forests | Tropical rainforest | Tropical forest | Tropical rainforest |
| | Tropical & subtropical dry broadleaf forests | Tropical woodland | | Sub-tropical forest |
| | | Monsoon or dry forest | | Tropical seasonal forest |
| | | Tropical thorn scrub and scrub woodland | | |
| | Tropical & subtropical coniferous forests | Montane tropical forest | | |
| | Mangroves | | | |
| **Temperate forest** | Temperate broadleaf & mixed forests | Broadleaved temperate evergreen forest | Warm temperate forest | Temperate broadleaved deciduous forest |
| | Mediterranean forests, woodlands & scrub | Semi-arid temperate woodland or scrub | | Warm temperate open woodland |
| | | | | Warm temperate mixed forest |
| | | | | Warm temperate broadleaved evergreen forest |
| | Temperate conifer forests | | Temperate forest | Cool temperate conifer forest |
| | | | | Cool temperate mixed forest |
| **Boreal forest** | Boreal forests/taiga | Open boreal woodlands | Boreal forest | Boreal/montane forest |
| | | Main Taiga | | Cold temperate/boreal open woodland |
| **Savanna and grassland** | Tropical & subtropical grasslands, savannas & shrublands | Tropical grassland | Savanna &dry woodland | Tropical savanna |
| | | Savanna | Grassland &dry scrubland | Tropical grassland |
| | Temperate grasslands, savannas & shrublands | Temperate steppe grassland | | Temperate grassland |
| | | Forest steppe | | |
| | | Dry steppe | | |
| | Flooded grasslands & savannas | | | |
| | Montane grasslands & shrublands | Alpine tundra | | |
| | | Montane Mosaic | | |
| | | Subalpine parkland | | |
| **Tundra** | Tundra | Tundra | Tundra | Tundra |
| | | Steppe-tundra | Dry tundra | |
| | | Polar and alpine desert | | |
| **Deserts** | Deserts & xeric shrublands | Tropical semi-desert | Desert | Desert |
| | | Tropical extreme desert | | Semi-desert |
| | | Temperate desert | | |
| | | Temperate semi-desert | | |
| **Not vegetated** | Lake | Lakes and open water | | |
| | Rock and ice | Ice sheet and other permanent ice | Ice | Ice |

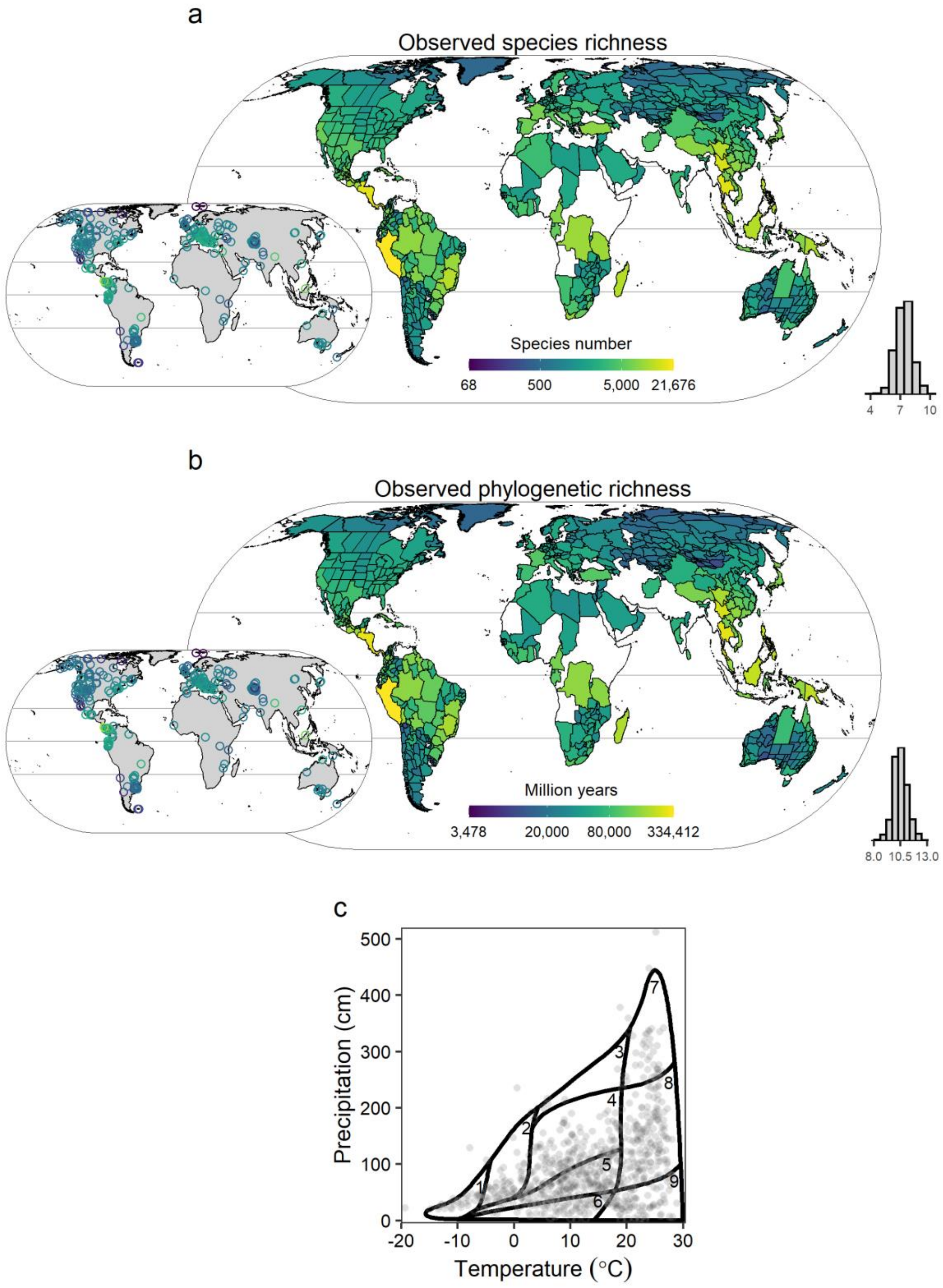


**Supplementary Fig. 1 | Observed species richness and phylogenetic richness of vascular plants for 830 geographic regions used to train the models.** a, species richness. b, phylogenetic richness. Embedded maps show observed species richness (a) and phylogenetic richness (b) for overlapping small regions and regions <10000 km$^2$. The histograms show the frequency distribution of log species richness (a) and log phylogenetic richness (b). c, Regions plotted onto Whittaker's scheme of biomes delineated based on annual mean temperature and annual precipitation[22]. Whittaker biomes are numbered as follows: 1 = tundra , 2 = boreal forest, 3 = temperate rainforest , 4 = temperate seasonal forest , 5 = woodland/shrubland , 6 = temperate grassland, 7 = tropical rainforest, 8 = tropical seasonal forest, 9 = subtropical desert.

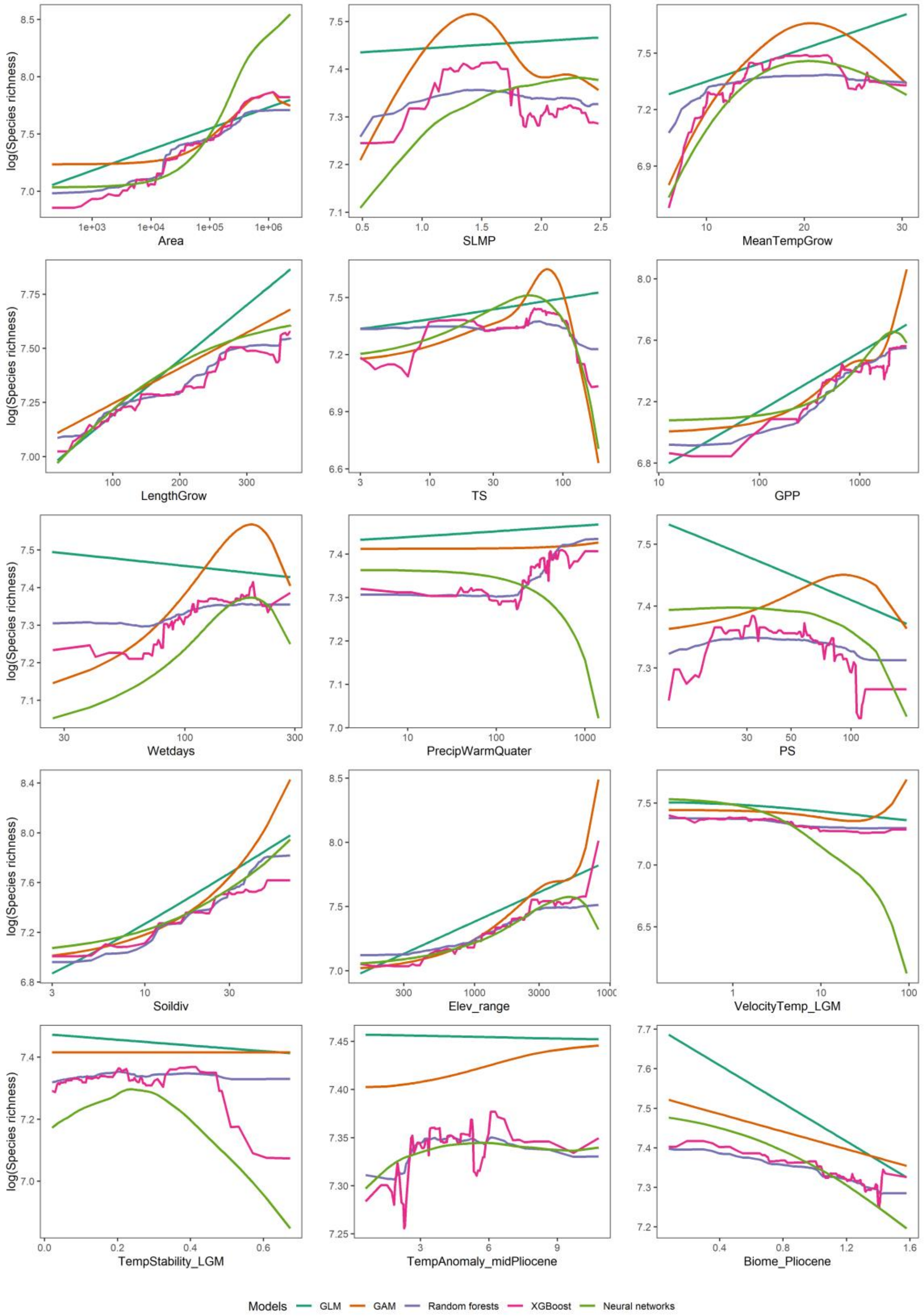


**Supplementary Fig. 2 | Estimated effects of predictor variables on species richness of vascular plants across five non-spatial models (partial dependence plots).** Five non-spatial models were fitted with 15 predictors (see Supplementary Table 1 for predictor abbreviations). Predictors shown here were used for both non-spatial and spatial models, and were selected to represent the major hypotheses related to plant diversity–environment relationships and filtered based on their contribution to model performance and collinearity.

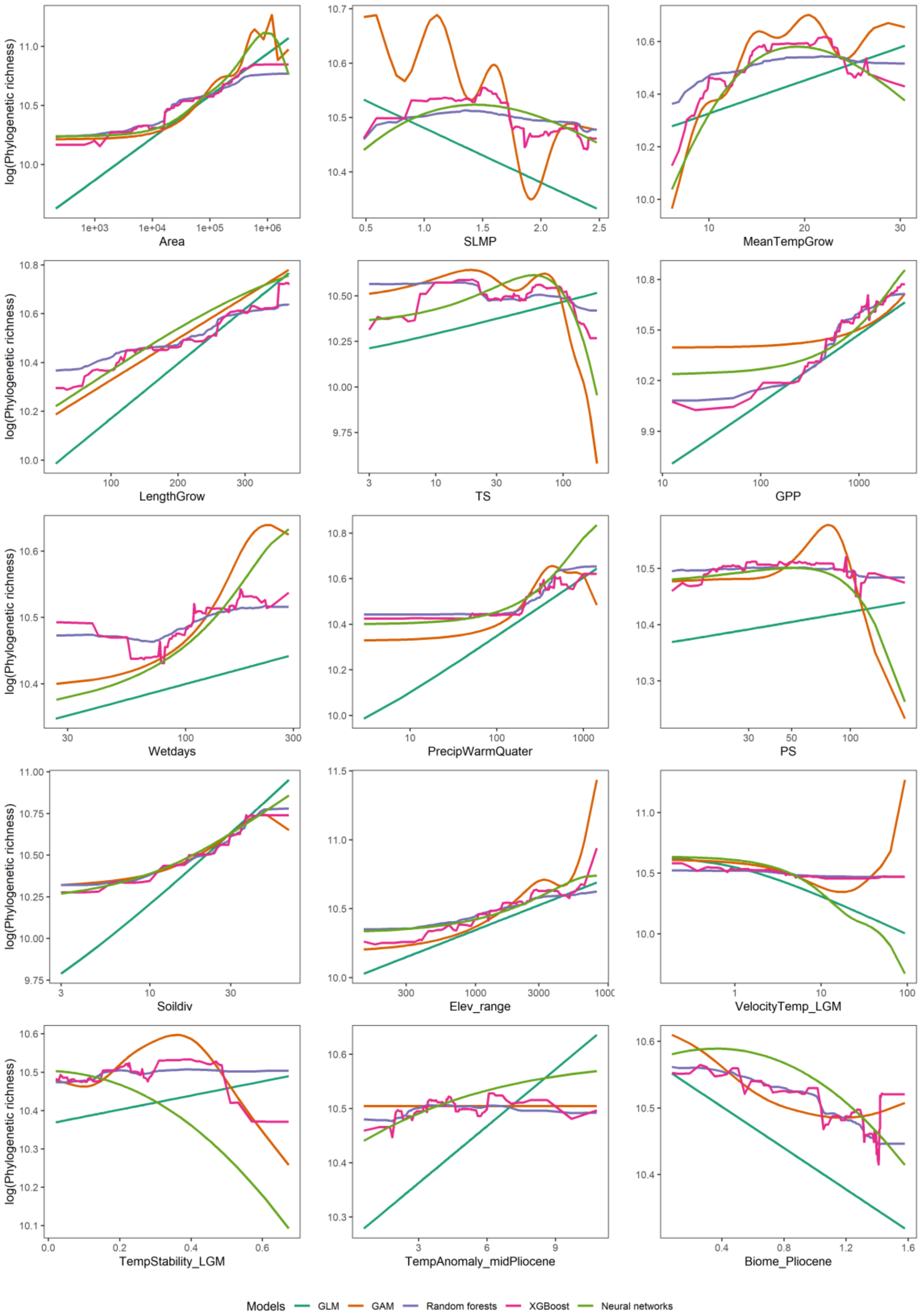


**Supplementary Fig. 3 | Estimated effects of predictor variables on phylogenetic richness of vascular plants across five non-spatial models (the partial dependence plots).** Five non-spatial models were fitted with 15 predictors (see Supplementary Table 1 for predictor abbreviations). Predictors shown here were used for both non-spatial and spatial models, and were selected to represent the major hypotheses related to plant diversity–environment relationships and filtered based on their contribution to model performance and collinearity.

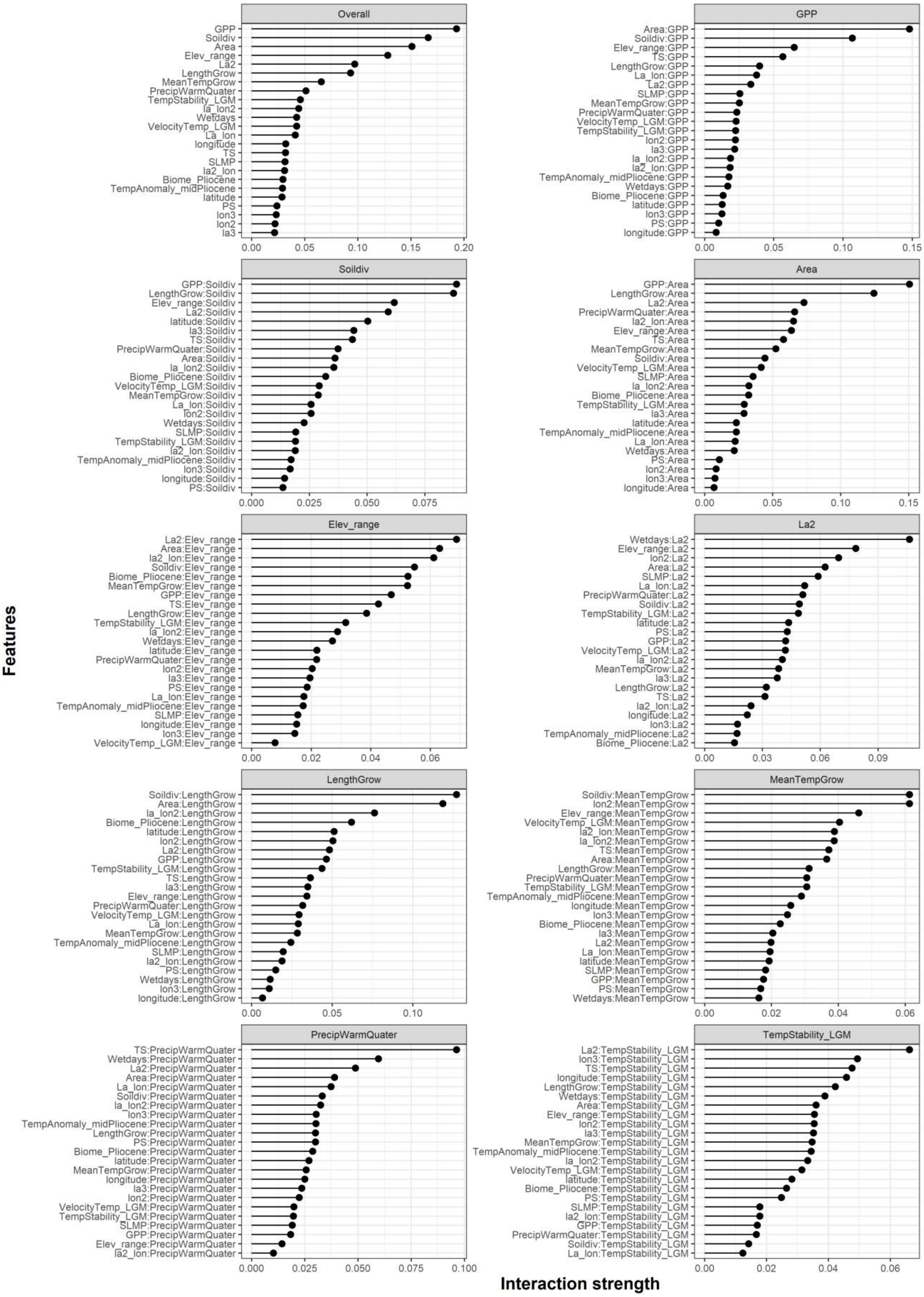


**Supplementary Fig. 4 | Interaction strength of each predictor variable for explaining species richness (Overall) in the spatial random forest model and two-way interaction strengths between the nine top-ranked covariates and all other covariates.** Terms of cubic polynomial trend surfaces (i.e. latitude (Y), centred longitude (X) as well as $X^2$, XY, $Y^2$, $X^3$, $X^2Y$, $XY^2$ and $Y^3$) are “La”, “Lon”, “La2”, “La_lon”, “lon2”, “la3”,”la2_lon”, “la_lon2”, and “lon3”, respectively. For other predictor abbreviations see Supplementary Table 1.

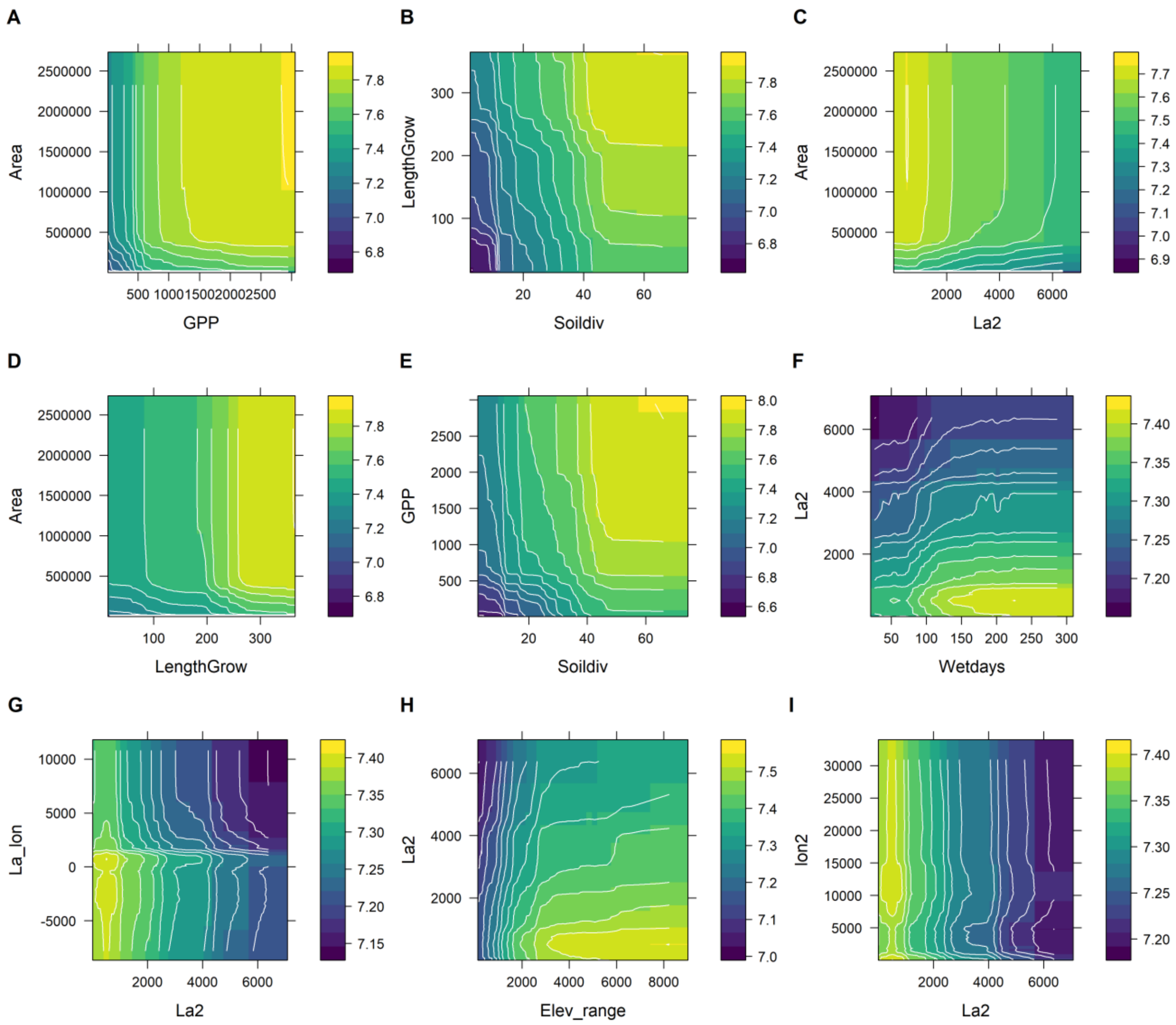


**Supplementary Fig. 5 | Estimated effects of the nine two-way interactions (two-predictors partial dependence plots) in the spatial random forest model for species richness.**

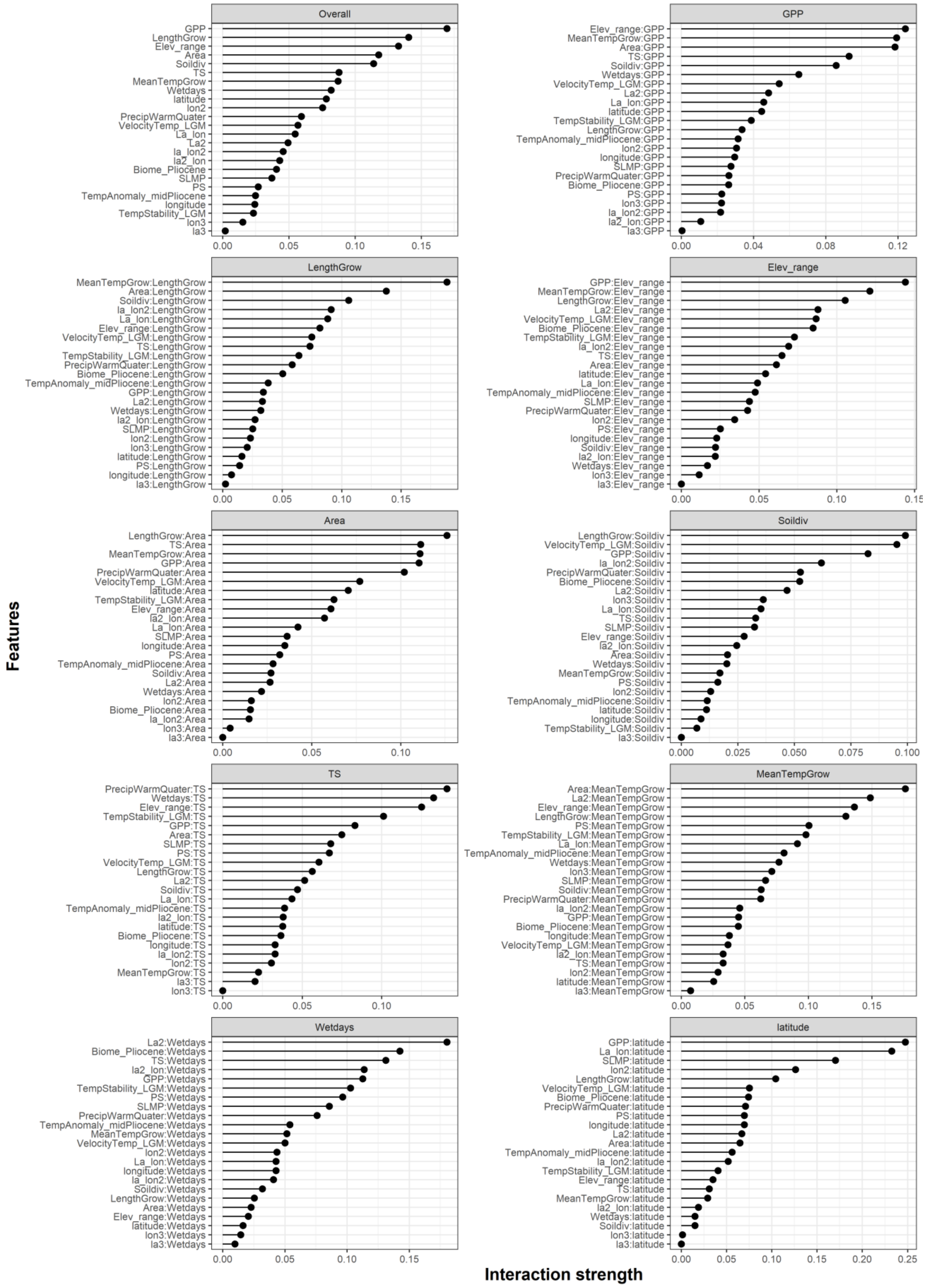


**Supplementary Fig. 6 | Interaction strength of each predictor variable for explaining species richness (Overall) in the spatial XGBoost model and two-way interaction strengths between the nine top-ranked covariates and all other covariates.**

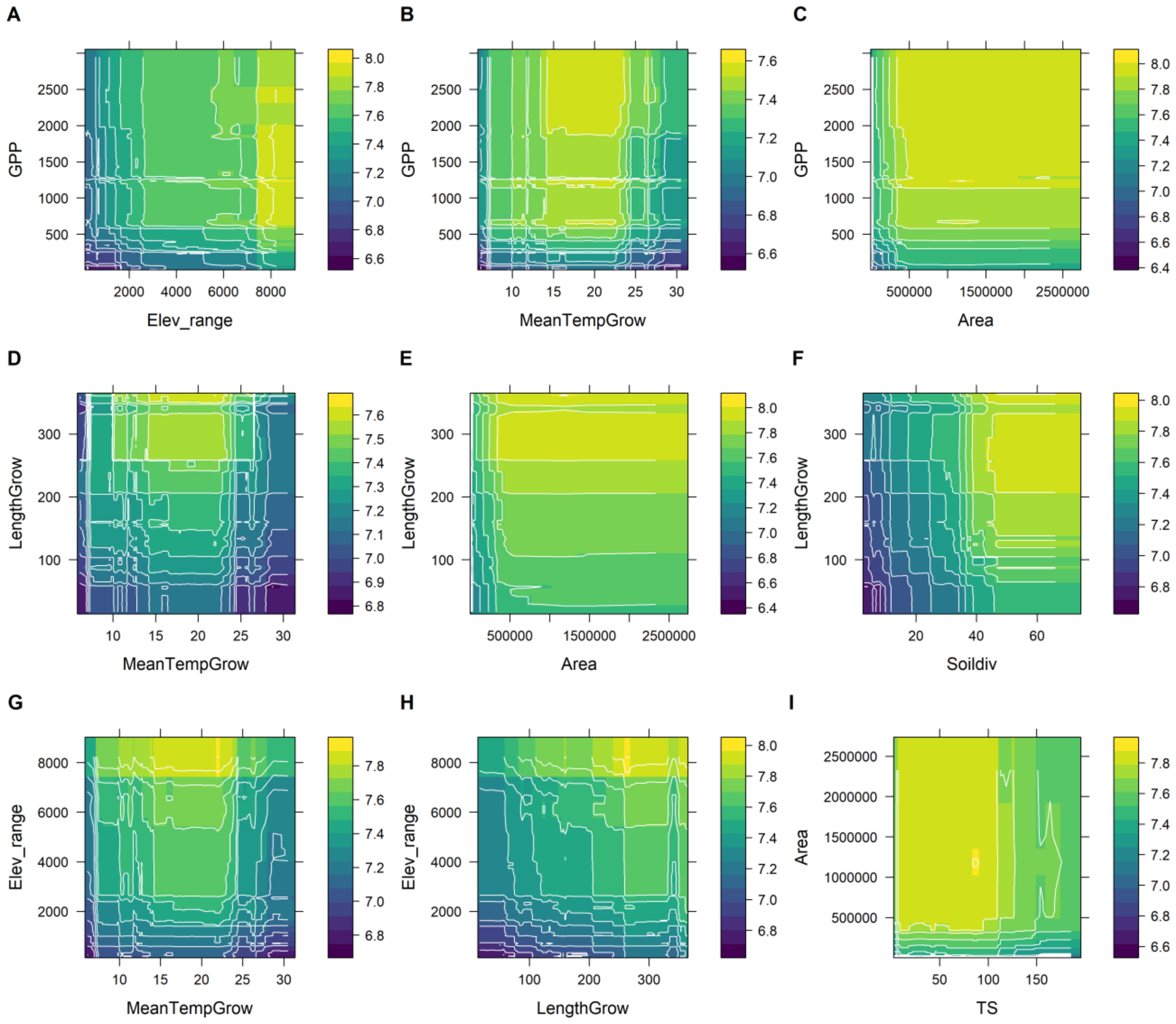


**Supplementary Fig. 7 | Estimated effects of the nine two-way interactions (two-predictors partial dependence plots) in the spatial XGBoost model for species richness.**

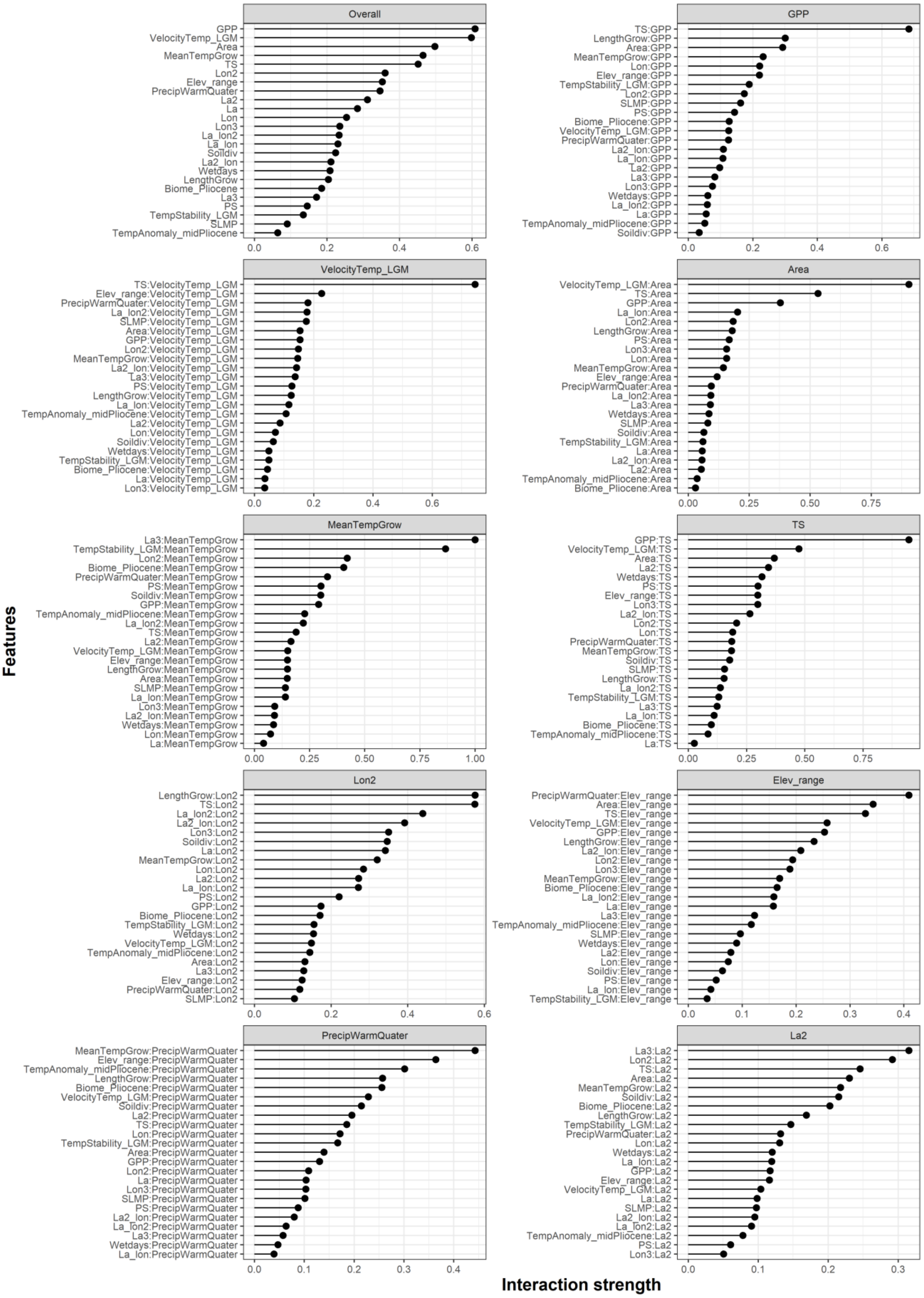

**Supplementary Fig. 8 | Interaction strength of each predictor variable for explaining species richness (Overall) in the spatial neural network model and two-way interaction strengths between the nine top-ranked covariates and all other covariates.**

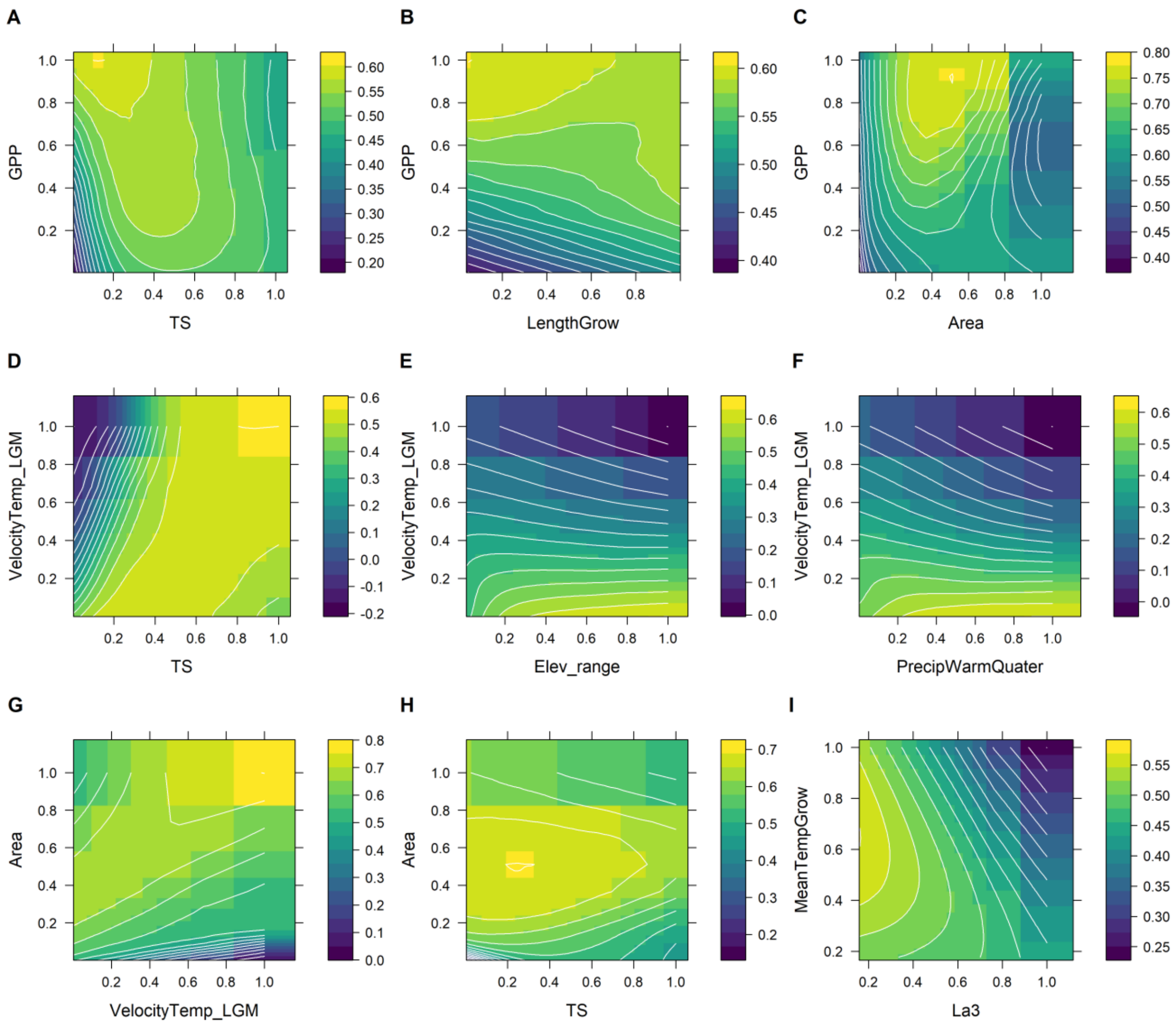


**Supplementary Fig. 9 | Estimated effects of the nine two-way interactions (two-predictors partial dependence plots) in the spatial neural network model for species richness.**

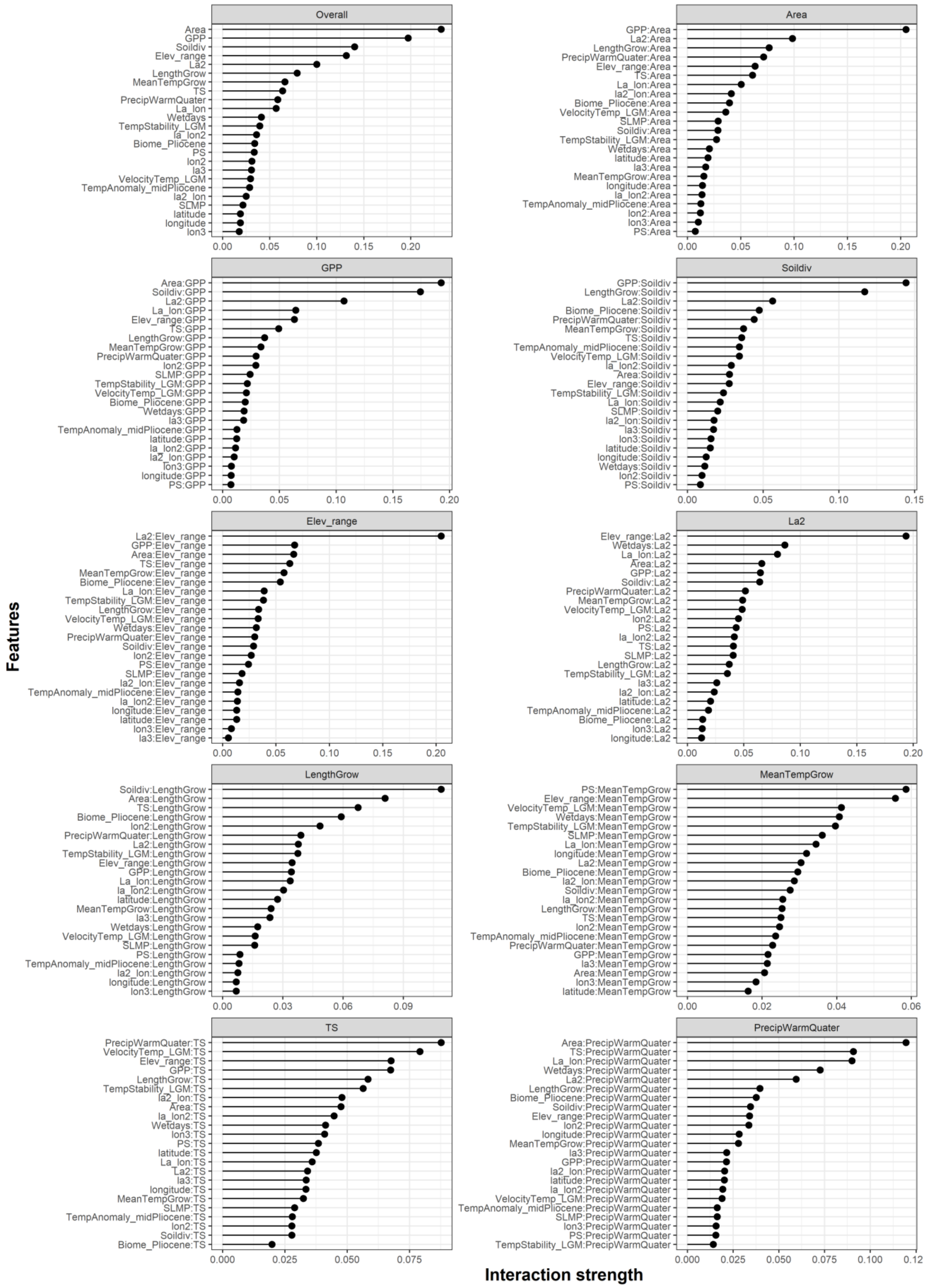

**Supplementary Fig. 10 | Interaction strength of each predictor variable for explaining phylogenetic richness (Overall) in the spatial random forest model and two-way interaction strengths between the nine top-ranked covariates and all other covariates.**

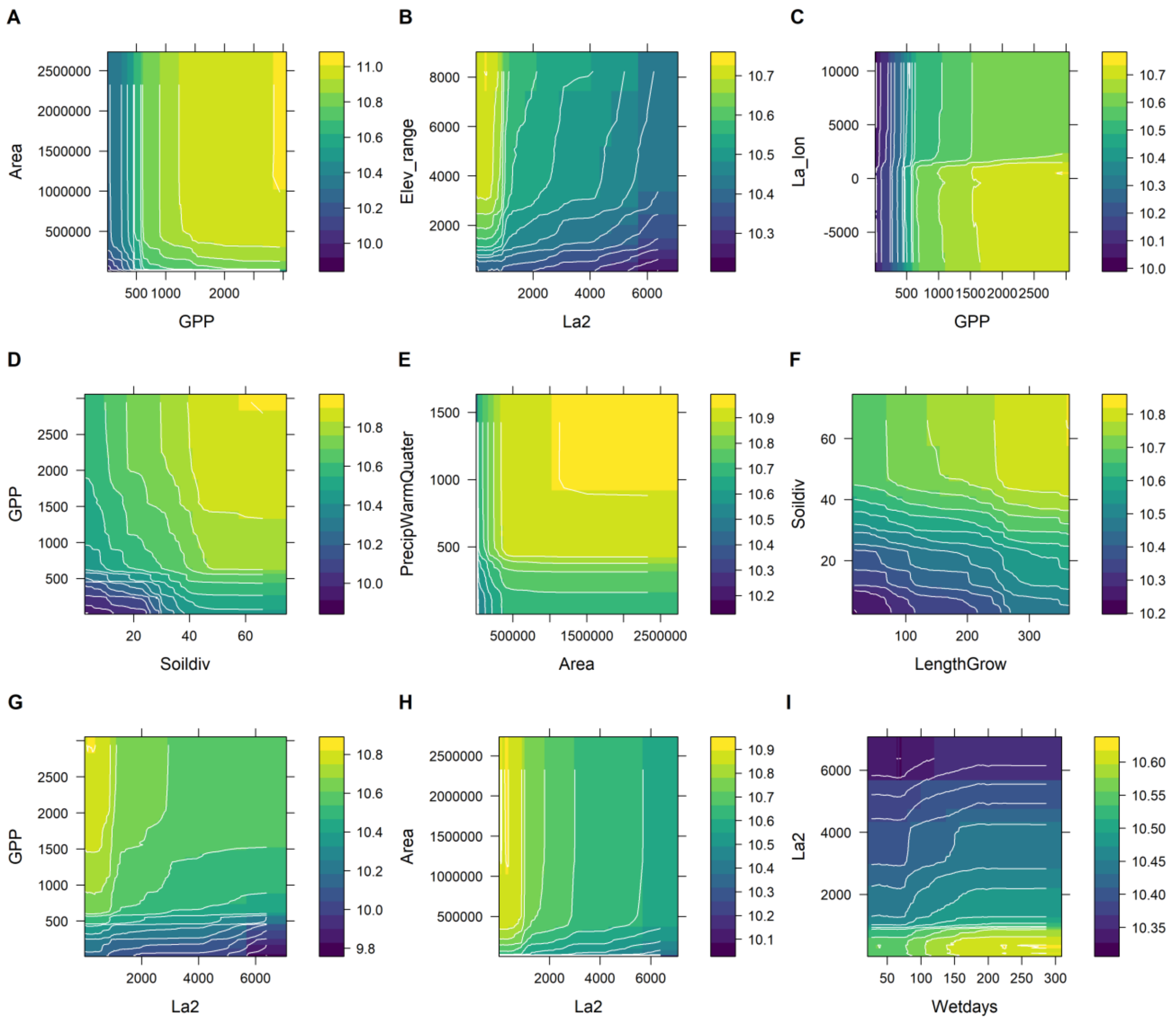


**Supplementary Fig. 11 | Estimated effects of the nine two-way interactions (two-predictors partial dependence plots) in the spatial random forest model for phylogenetic richness.**

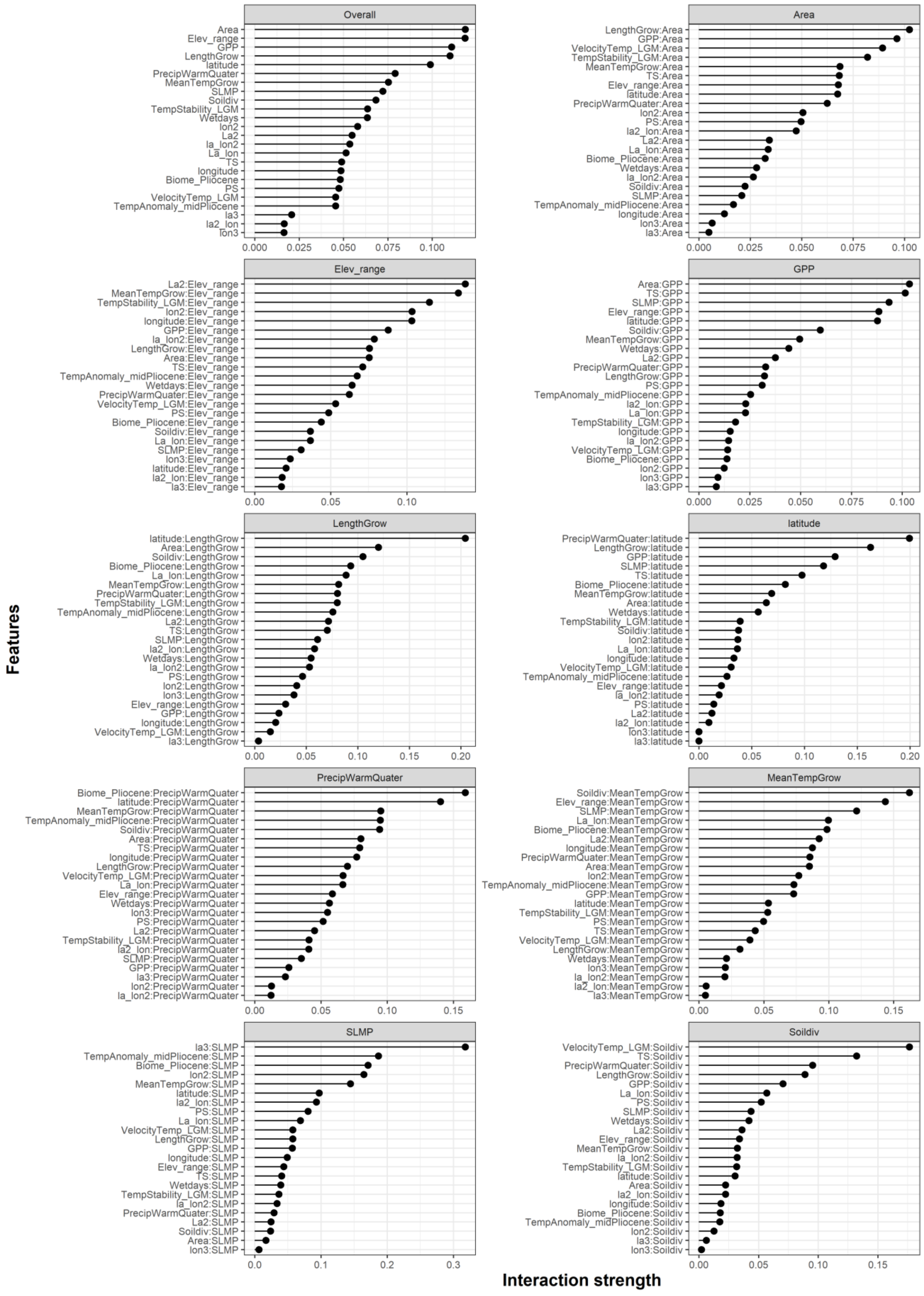


**Supplementary Fig. 12 | Interaction strength of each predictor variable for explaining phylogenetic richness (Overall) in the spatial XGBoost model and two-way interaction strengths between the nine top-ranked covariates and all other covariates.**

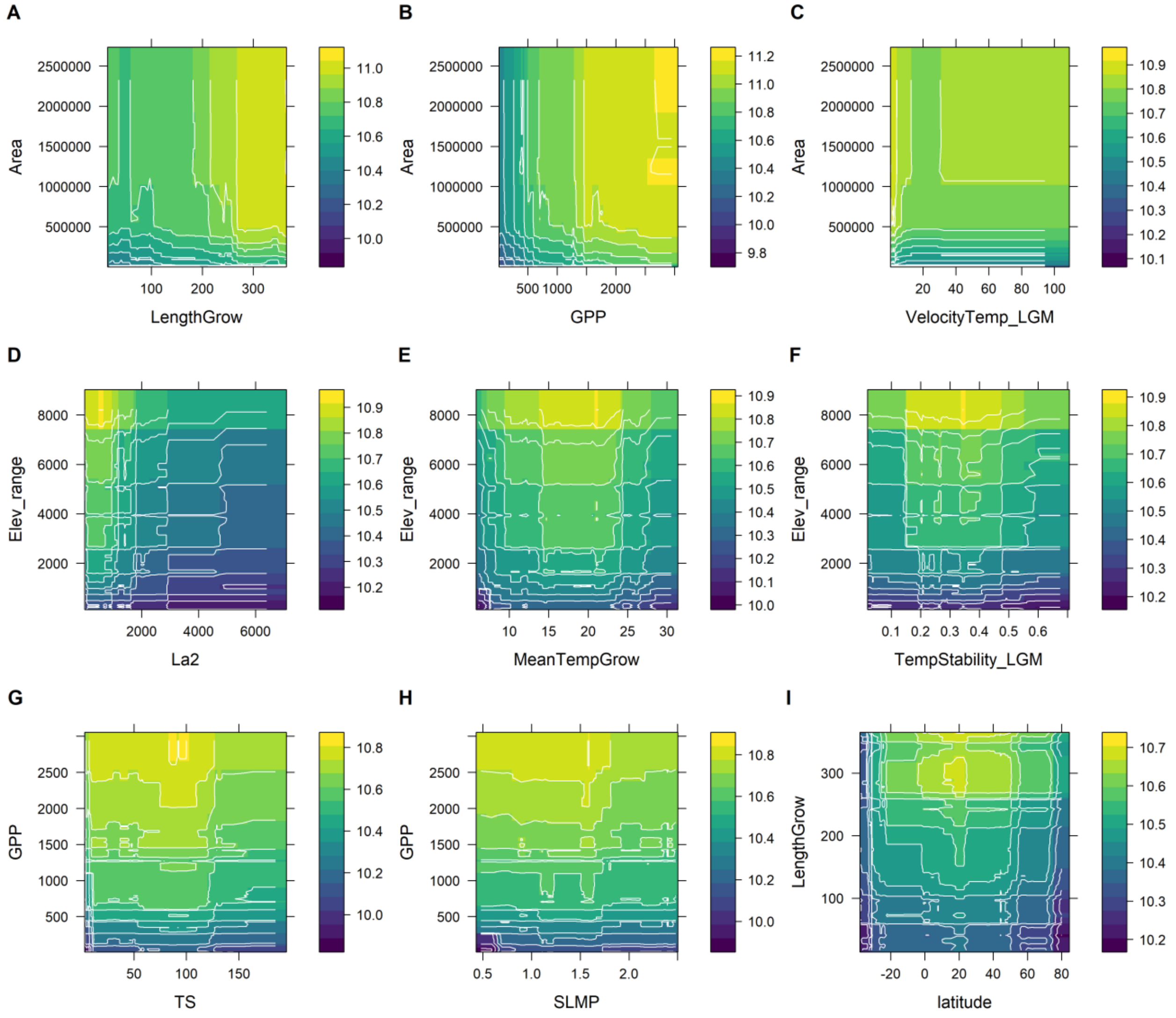


**Supplementary Fig. 13 | Estimated effects of the nine two-way interactions (two-predictors partial dependence plots) in the spatial XGBoost model for phylogenetic richness.**

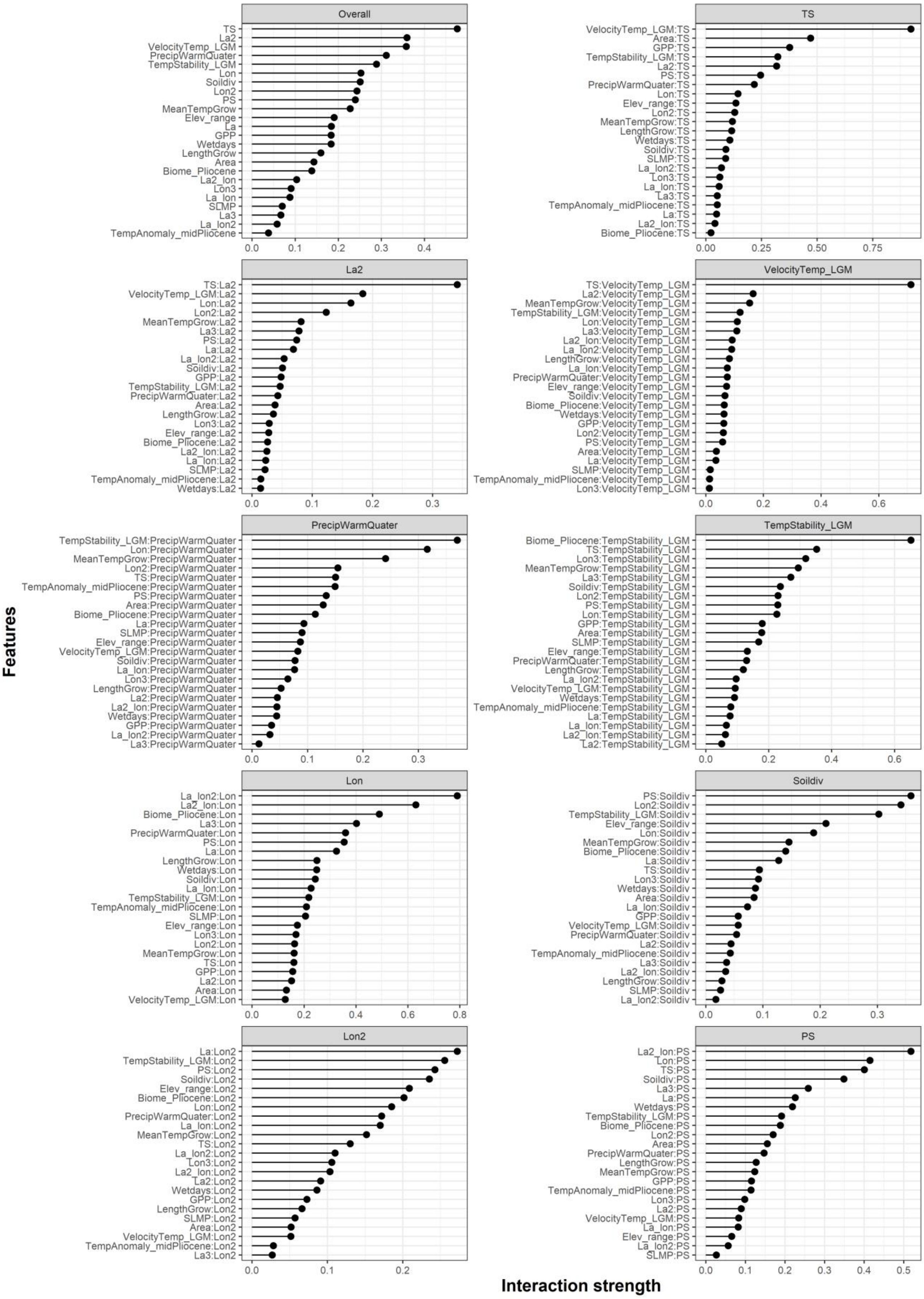


**Supplementary Fig. 14 | Interaction strength of each predictor variable for explaining phylogenetic richness (Overall) in the spatial neural network model and two-way interaction strengths between the nine top-ranked covariates and all other covariates.**

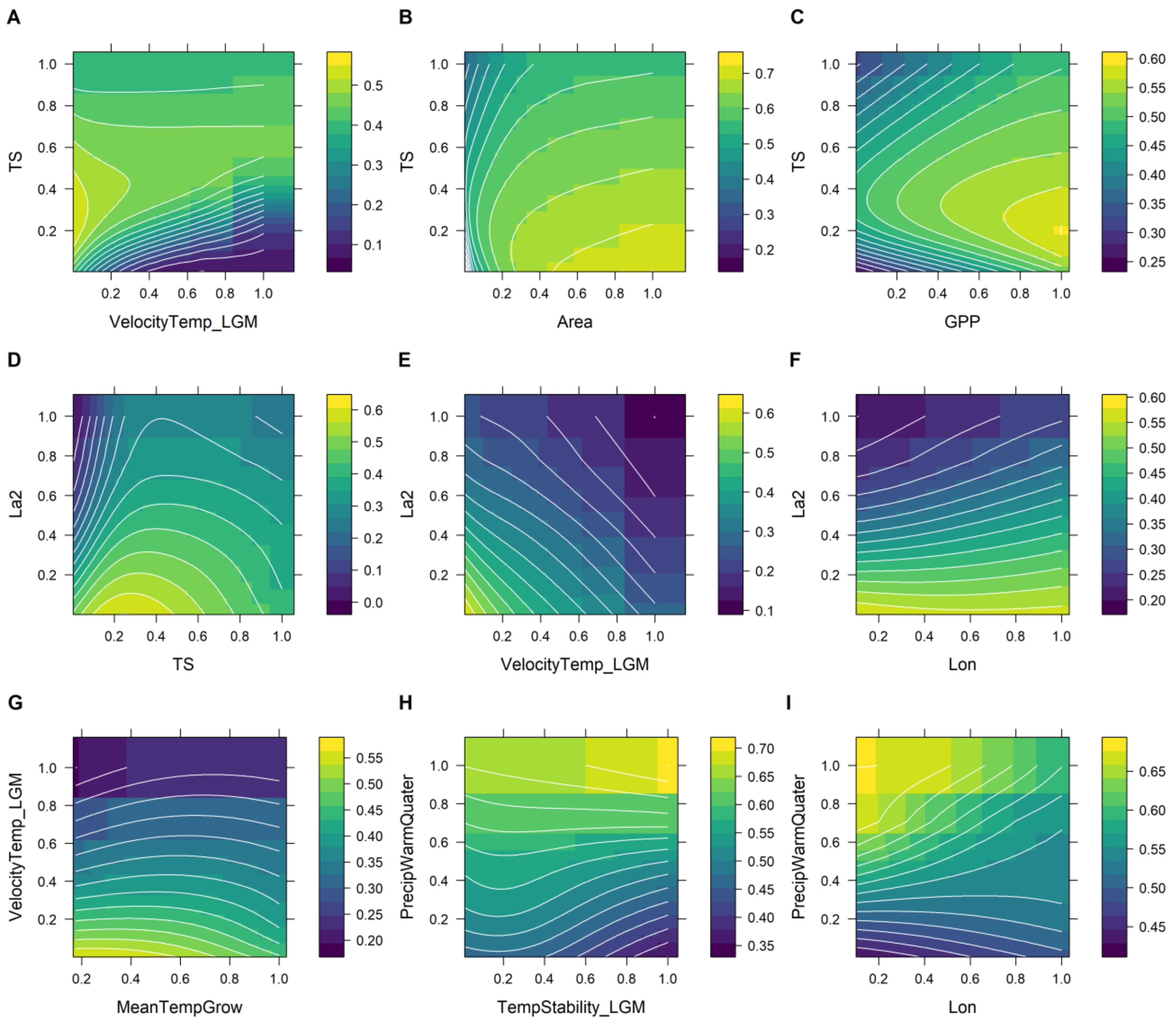


**Supplementary Fig. 15 | Estimated effects of the nine two-way interactions (two-predictors partial dependence plots) in the spatial neural network model for phylogenetic richness.**

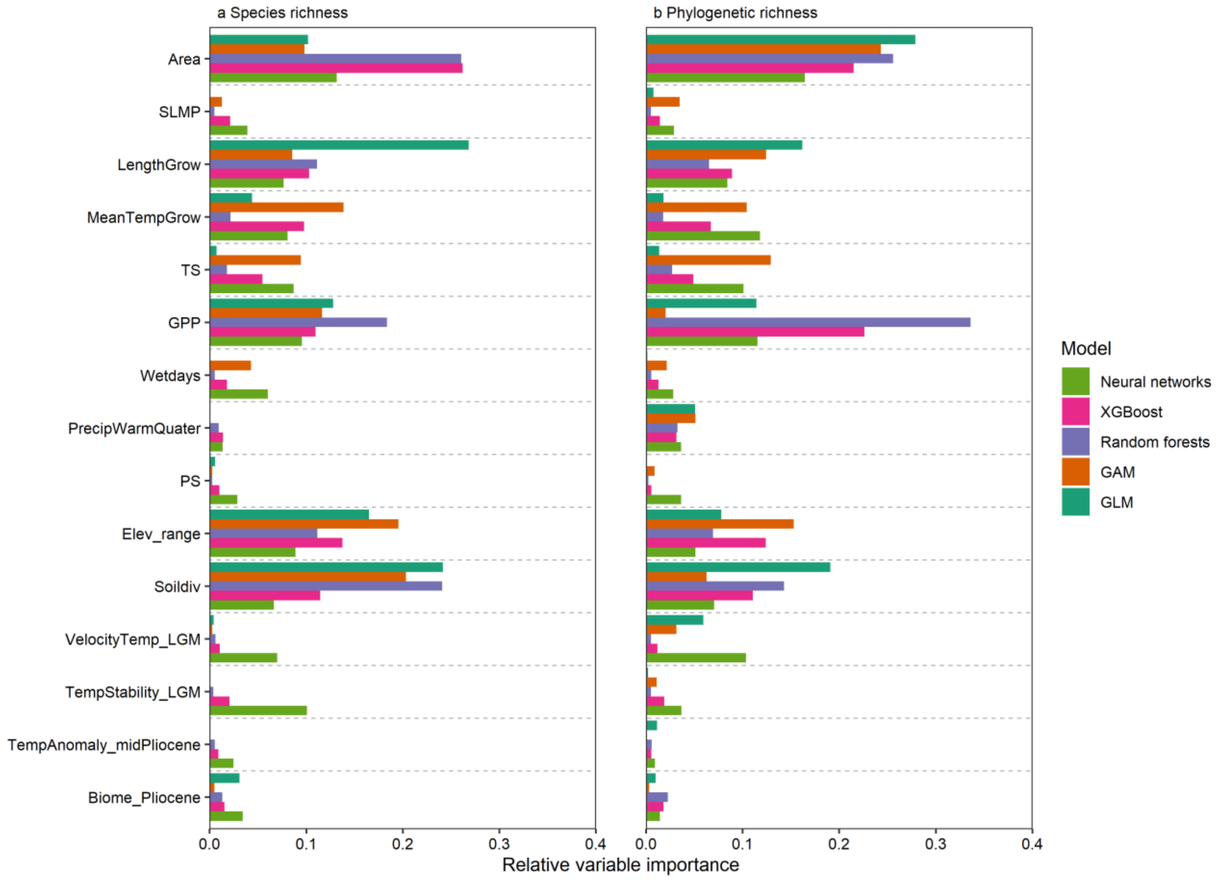


**Supplementary Fig. 16 | Relative importance of environmental variables explaining global pattern of vascular plant diversity across five non-spatial models.** The models were fitted including 15 predictors representing geography, climate, environmental heterogeneity and past environmental conditions (see Supplementary Table 1 for predictor abbreviations). a, Importance of predictor variables for species richness; b, Importance of predictor variables for phylogenetic richness.

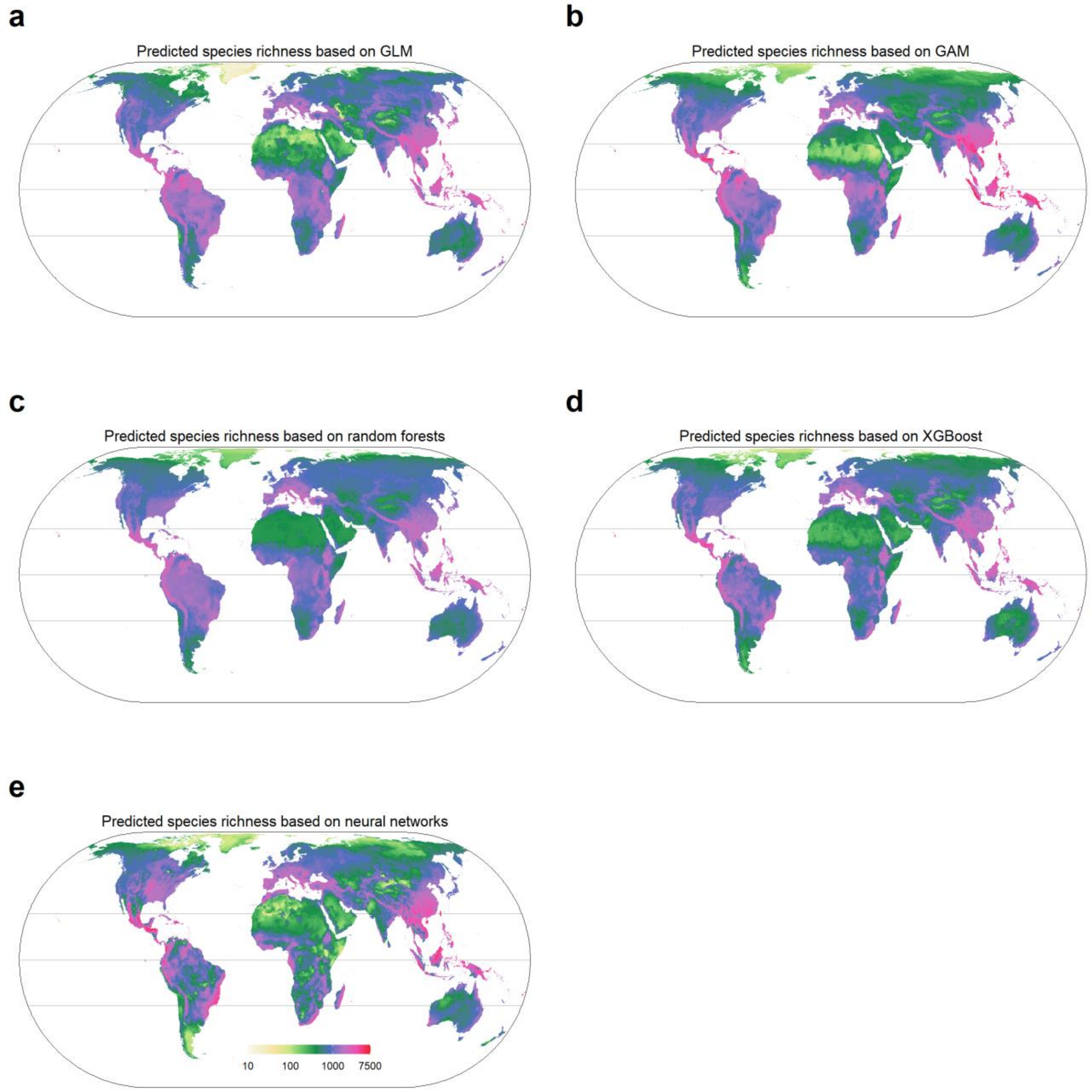


**Supplementary Fig. 17 | Species richness of vascular plants predicted across an equal area grid of 7,774 km$^2$ hexagons based on different models** (i.e. spatial models using machine learning methods and GAM, and a non-spatial GLM with interactions)**.** The same $\log_{10}$ scale colour gradient is used in all maps. For comparisons across all spatial and non-spatial models and data download, see https://gift.uni-goettingen.de/shiny/predictions/. Projection: Eckert IV.

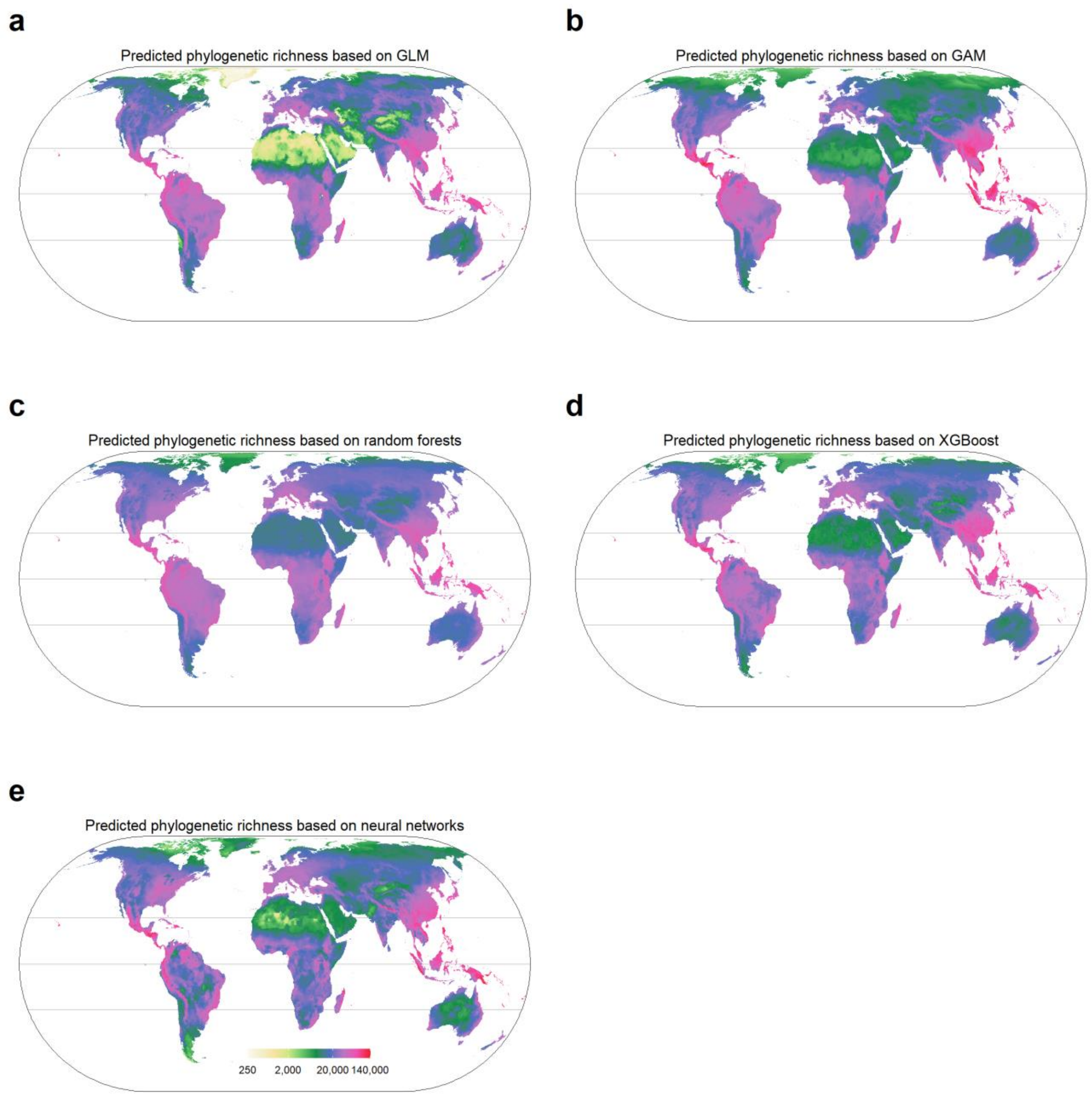


**Supplementary Fig. 18 | Phylogenetic richness of vascular plants predicted across an equal area grid of 7,774 km$^2$ hexagons based on different models** (i.e. spatial models using machine learning methods and GAM, and a non-spatial GLM with interactions)**.** The same $\log_{10}$ scale colour gradient is used in all maps. For comparisons across all spatial and non-spatial models and data download, see https://gift.uni-goettingen.de/shiny/predictions/. Projection: Eckert IV.

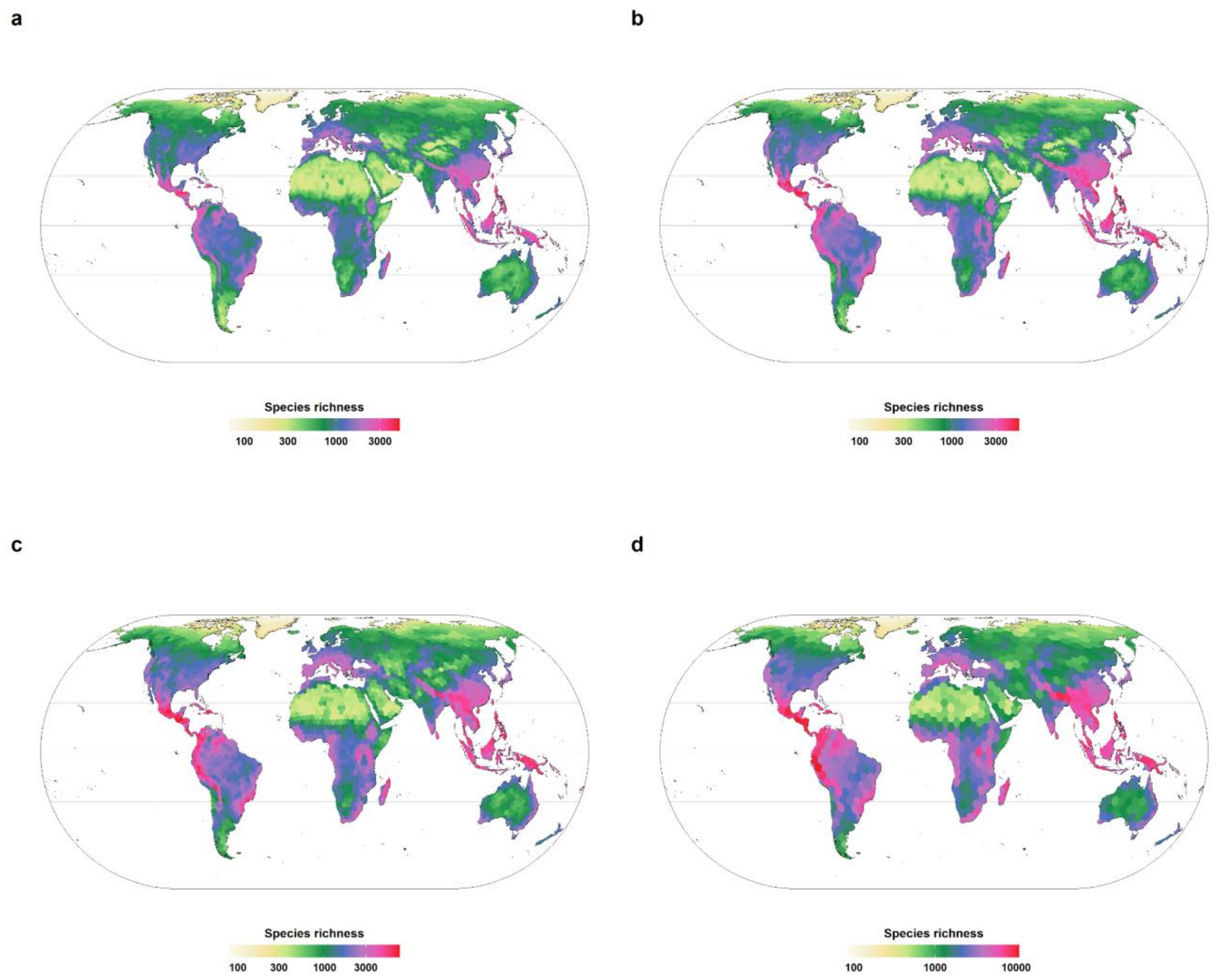


**Supplementary Fig. 19 | Species richness of vascular plants based on ensemble predictions across different grid sizes** (i.e. spatial models using machine learning methods and GAM, and a non-spatial GLM with interactions). Grid sizes used for maps are: a, 7774 km²; b, 23322 km²; c, 69967 km²; d, 209903 km². For comparisons across all spatial and non-spatial models and data download, see https://gift.uni-goettingen.de/shiny/predictions/. Projection: Eckert IV.

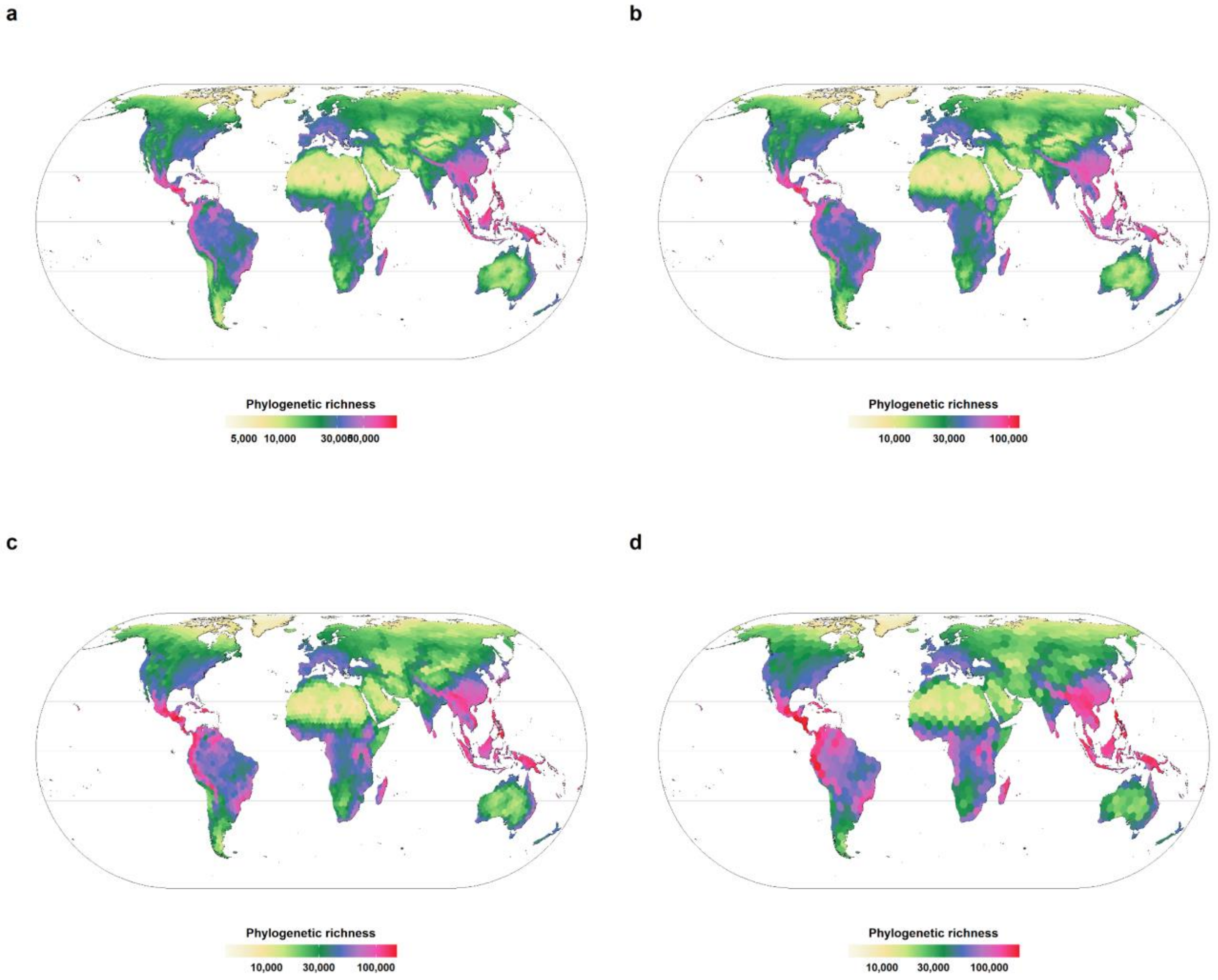


**Supplementary Fig. 20 | Phylogenetic richness of vascular plants based on ensemble predictions across different grid sizes** (i.e. spatial models using machine learning methods and GAM, and a non-spatial GLM with interactions). Grid sizes used for maps are: a, 7774 km²; b, 23322 km²; c, 69967 km²; d, 209903 km². For comparisons across all spatial and non-spatial models and data download, see https://gift.uni-goettingen.de/shiny/predictions/. Projection: Eckert IV.

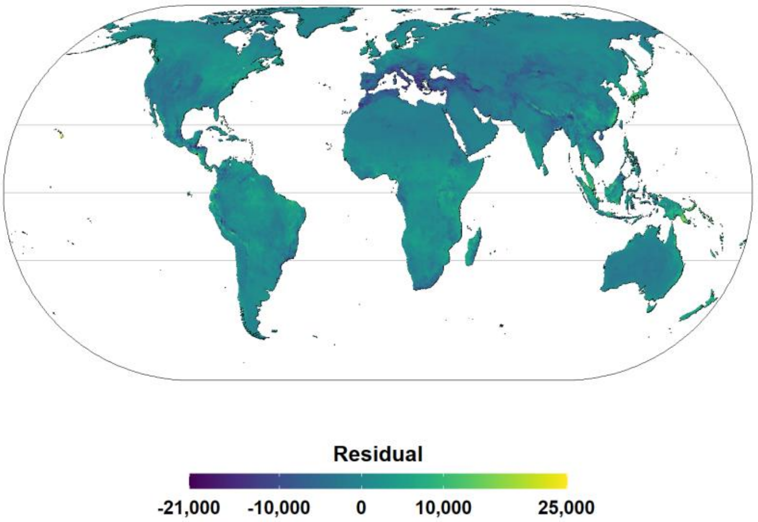


**Supplementary Fig. 21 | Residuals (deviation) from the linear regression between species richness and phylogenetic richness based on Ensemble predictions** (phylogenetic richness = 22.1 * species richness, R2= 0.947, p < 0.0001). Negative residuals indicate lower phylogenetic richness than expected based on species richness. Projection: Eckert IV.

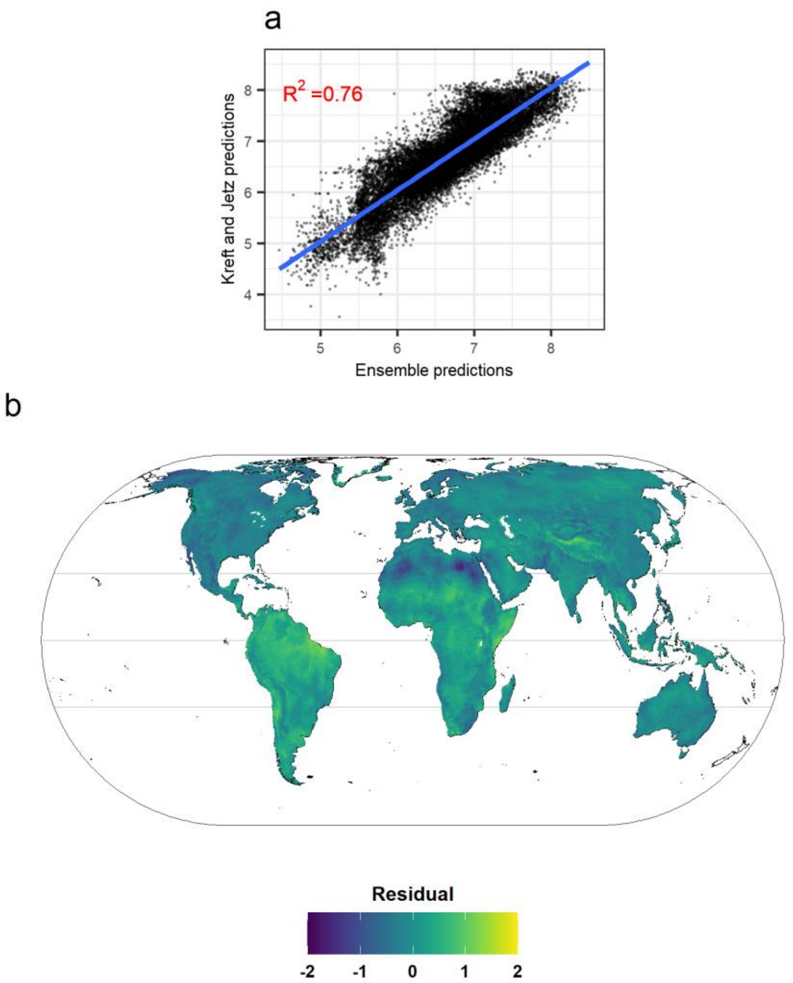


**Supplementary Fig. 22 | Comparison between vascular plant species richness based on Ensemble predictions produced in the scope of this paper (SR.Ensemble) and species richness extracted from Kreft and Jetz[20] (SR.Kreft)** (a, SR.Kreft = 1.01 SR.Ensemble, $R^2$= 0.76, p < 0.0001). Species richness was log-transformed. b, global patterns of residuals from the linear regression between species richness based on the ensemble predictions and species richness extracted from Kreft and Jetz's[20]. Positive residuals indicate higher values of SR.Kreft. Projection: Eckert IV.

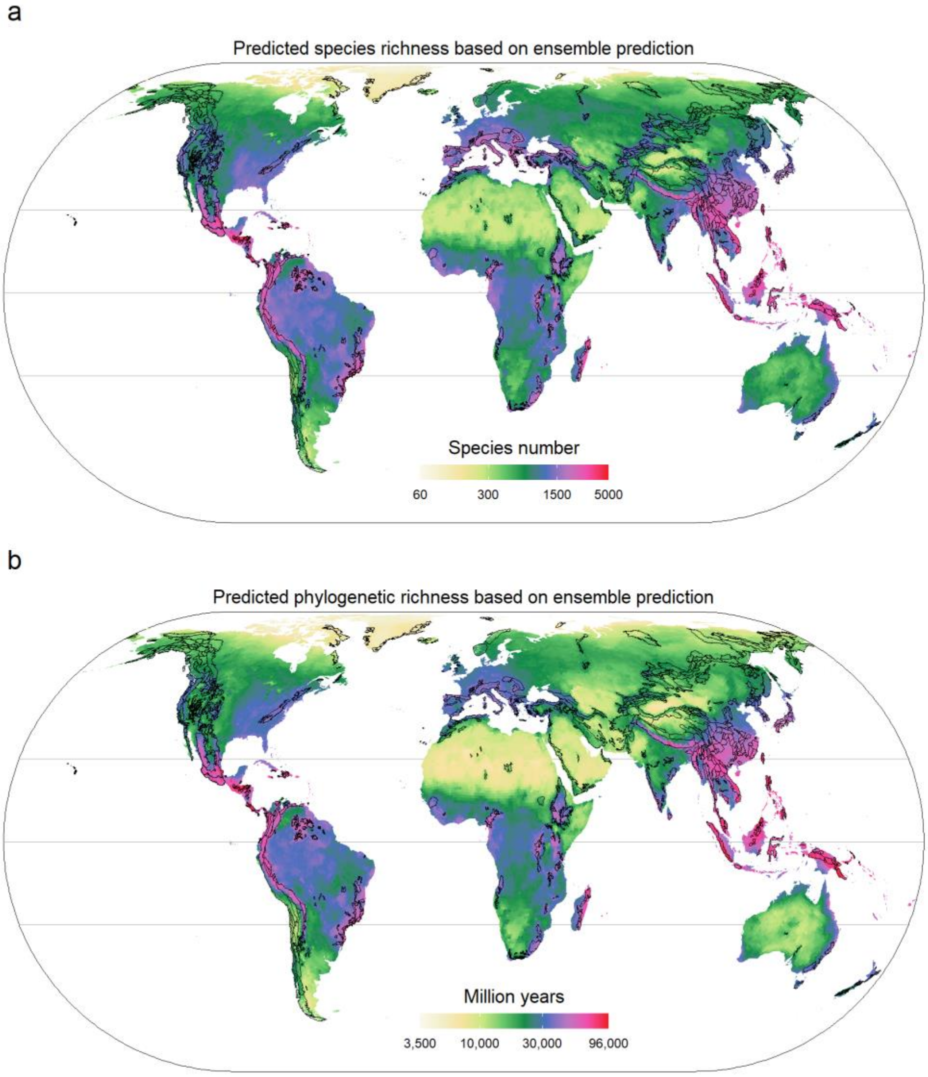


**Supplementary Fig. 23 | Vascular plant diversity based on ensemble predictions across an equal area grid of 7774 km$^2$ hexagons and mountain regions.** Black lines delineate mountainous regions worldwide based on Körner et al.[23]. Projection: Eckert IV.

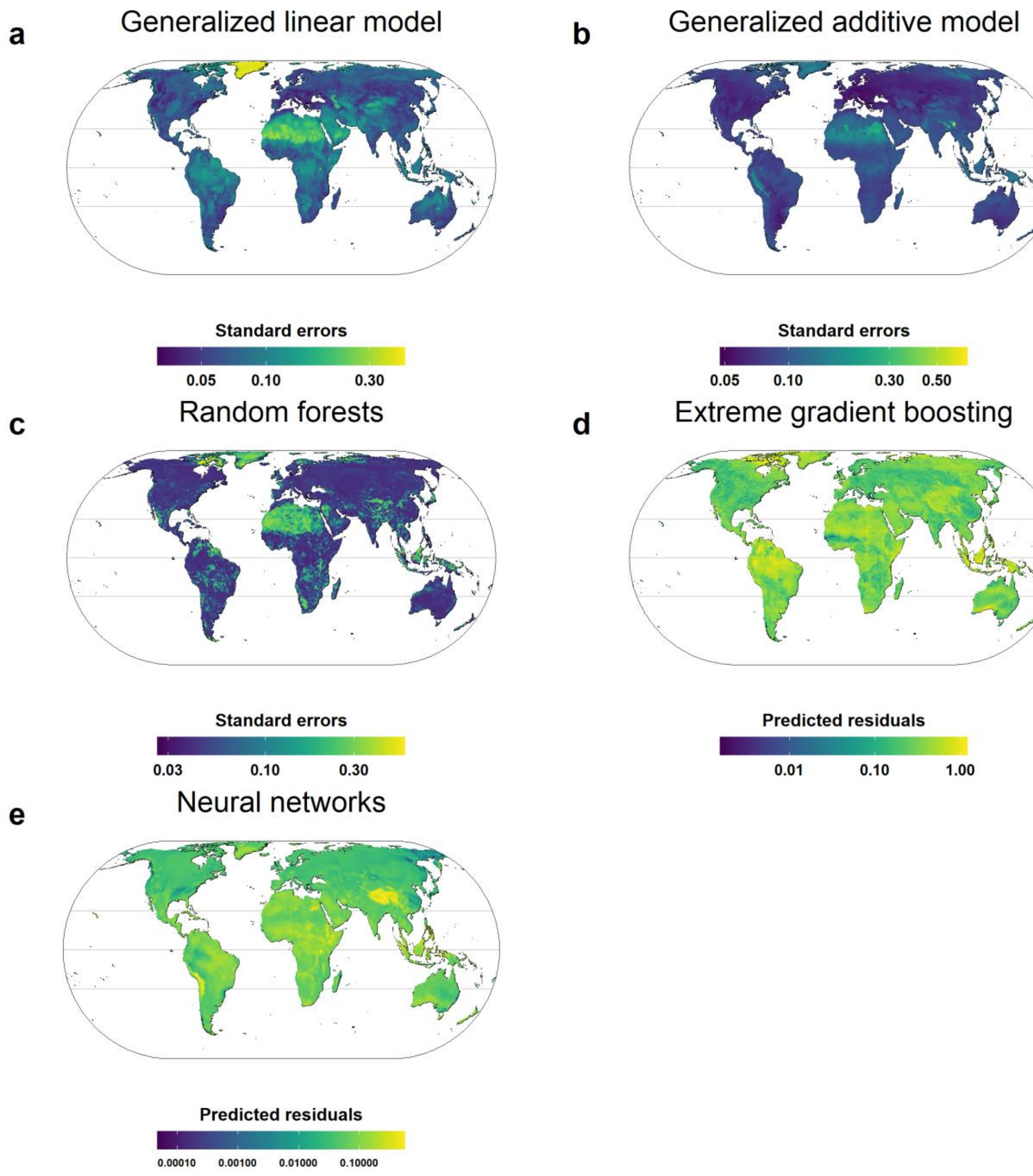


**Supplementary Fig. 24 | Uncertainty in predicted species richness from the five models used for the ensemble predictions** (i.e. spatial models using machine learning methods and GAM, and a non-spatial GLM with interactions). Prediction variation is measured as standard errors of predicted values in GAM, GLM and random forests and predicted residuals in XGBoost and neural networks based on models fitting the relationship between residuals of trained models and predictors from the raw datasets. For comparisons across all spatial and non-spatial models and data download, see https://gift.uni-goettingen.de/shiny/predictions/. Projection: Eckert IV.

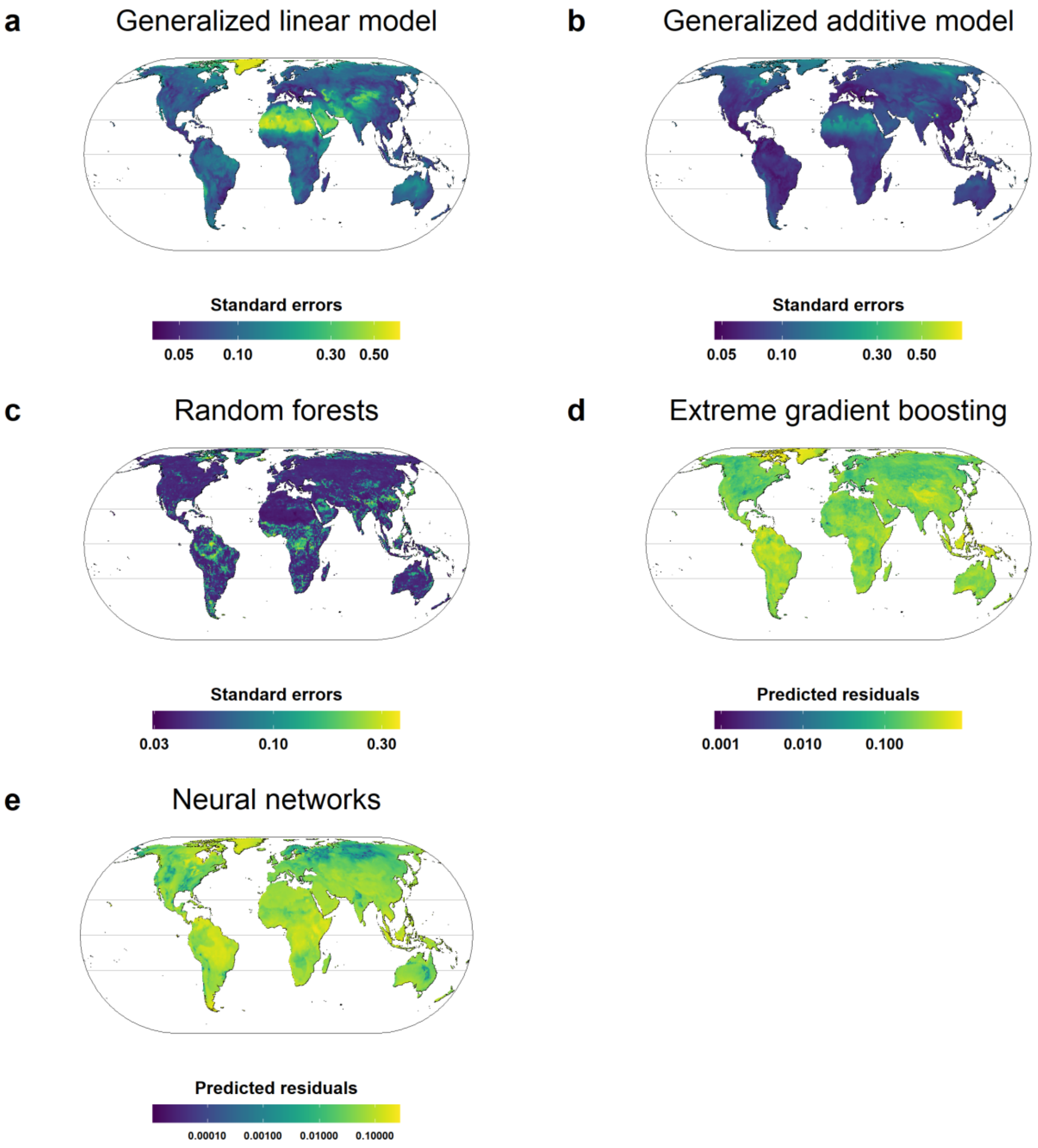


**Supplementary Fig. 25 | Uncertainty in predicted phylogenetic richness from the five models used for the ensemble predictions** (i.e. spatial models using machine learning methods and GAM, and a non-spatial GLM with interactions)**.** Prediction variation is measured as standard errors of predicted values in GAM, GLM and random forests and predicted residuals in XGBoost and neural networks based on models fitting the relationship between the residuals of trained models and predictors from the raw datasets. For comparisons across all spatial and non-spatial models and data download, see https://gift.uni-goettingen.de/shiny/predictions/. Projection: Eckert IV.

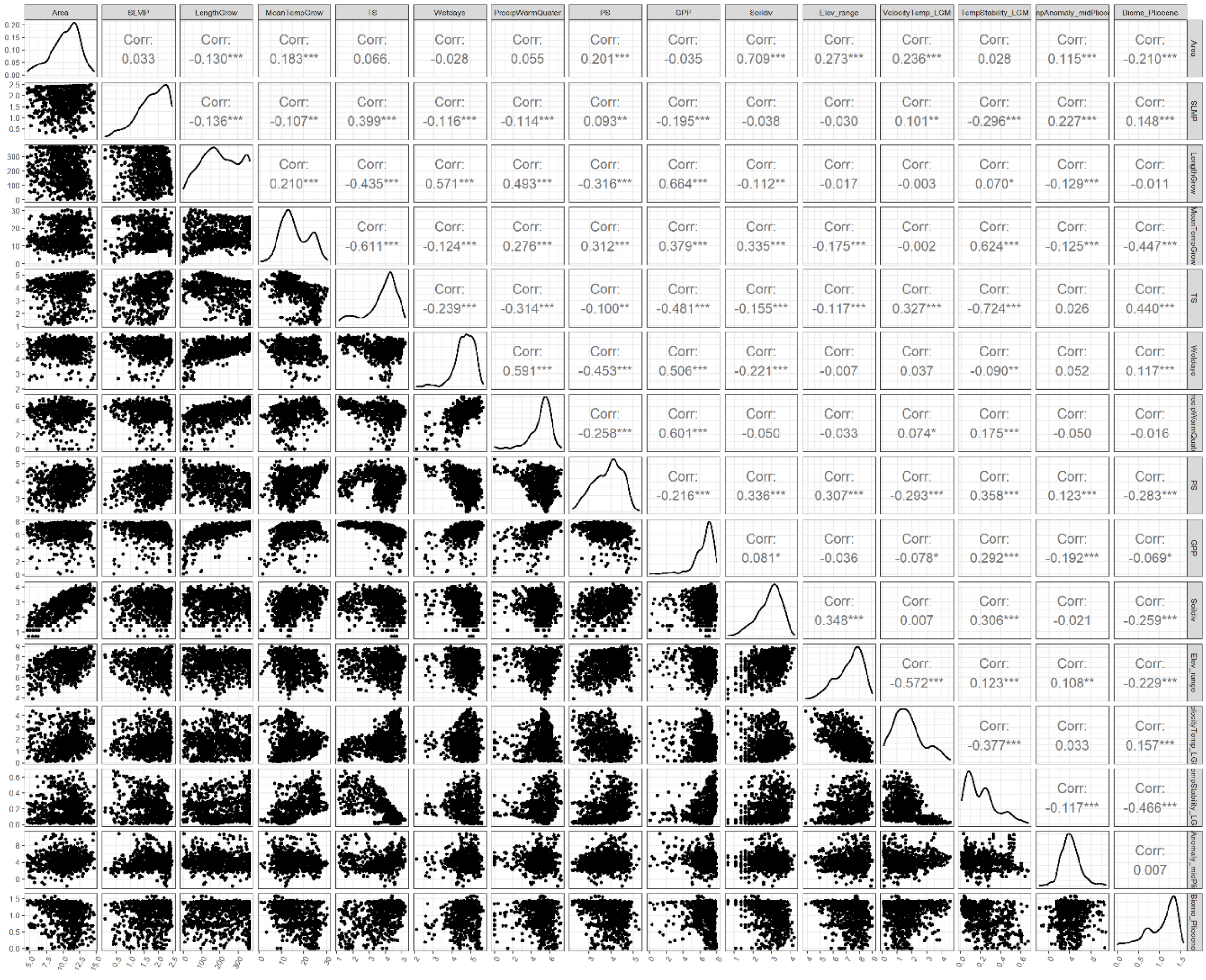


**Supplementary Fig. 26 | Correlations among all predictors and their density distributions.** Numbers are Pearson correlation coefficients. Some predictors (i.e. Area, TS, Wetdays, PrecipWarmQuater, PS, GPP, Soildiv, Elev_range, VelocityTemp_LGM; See Supplementary Table 1 for predictor abbreviations) are shown in log-scale as they were log-transformed for GLMs owing to their skewed distributions.

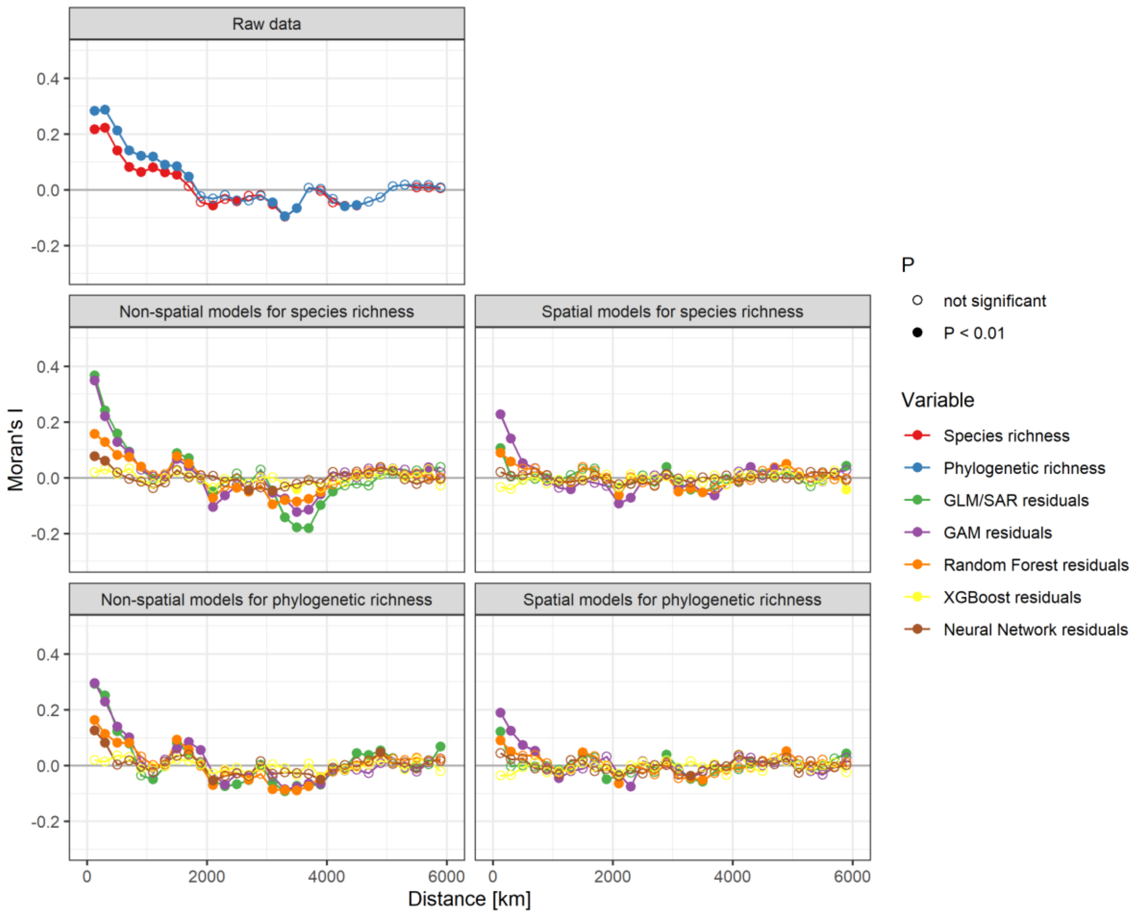


**Supplementary Fig. 27 | Spatial correlograms of raw diversity data, and residuals from non-spatial and spatial models, respectively, fitted for species richness and phylogenetic richness.** Full symbols indicate a significant Moran's I correlation at a given lag distance (P < 0.01).

**Supplementary references 1 | References used in the supplementary materials.**


1. Weigelt, P., König, C. & Kreft, H. GIFT – A global inventory of floras and traits for macroecology and biogeography. *J. Biogeogr.* **47**, 16–43 (2020).
2. Weigelt, P., Jetz, W. & Kreft, H. Bioclimatic and physical characterization of the world's islands. *Proc. Natl. Acad. Sci. USA* **110**, 15307–15312 (2013).
3. Karger, D. N. *et al.* Climatologies at high resolution for the earth's land surface areas. *Sci. Data* **4**, 170122 (2017).
4. Zomer, R. J., Trabucco, A., Bossio, D. A. & Verchot, L. V. Climate change mitigation: A spatial analysis of global land suitability for clean development mechanism afforestation and reforestation. *Agric. Ecosyst. Environ.* **126**, 67–80 (2008).
5. Karger, D. N. *et al.* Why tree lines are lower on islands—Climatic and biogeographic effects hold the answer. *Glob. Ecol. Biogeogr.* **28**, 839–850 (2019).
6. New, M., Lister, D., Hulme, M. & Makin, I. A high-resolution data set of surface climate over global land areas. *Clim. Res.* **21**, 1–25 (2002).
7. Zhao, M. & Running, S. W. Drought-induced reduction in global terrestrial net primary production from 2000 through 2009. *Science* **329**, 940–943 (2010).
8. Danielson, J. J. & Gesch, D. B. *Global multi-resolution terrain elevation data 2010 (GMTED2010).* https://pubs.er.usgs.gov/publication/ofr20111073 (2011).
9. Hengl, T. *et al.* SoilGrids250m: Global gridded soil information based on machine learning. *PLoS One* **12**, e0169748 (2017).
10. Tuanmu, M.-N. & Jetz, W. A global, remote sensing-based characterization of terrestrial habitat heterogeneity for biodiversity and ecosystem modelling. *Glob. Ecol. Biogeogr.* **24**, 1329–1339 (2015).
11. Hijmans, R. J., Cameron, S. E., Parra, J. L., Jones, P. G. & Jarvis, A. Very high resolution interpolated climate surfaces for global land areas. *Int. J. Climatol.* **25**, 1965–1978 (2005).
12. Owens, H. L. & Guralnick, R. climateStability: an R package to estimate climate stability from time-slice climatologies. *Biodivers. Inform.* **14**, 8–13 (2019).
13. Brown, J. L., Hill, D. J., Dolan, A. M., Carnaval, A. C. & Haywood, A. M. PaleoClim, high spatial resolution paleoclimate surfaces for global land areas. *Sci. Data* **5**, 1–9 (2018).
14. Hill, D. J. The non-analogue nature of Pliocene temperature gradients. *Earth Planet. Sci. Lett.* **425**, 232–241 (2015).
15. Dowsett, H. *et al.* The PRISM4 (mid-Piacenzian) paleoenvironmental reconstruction. *Clim. Past* **12**, 1519–1538 (2016).
16. Henrot, A.-J. *et al.* Effects of $CO^2$, continental distribution, topography and vegetation changes on the climate at the Middle Miocene: a model study. *Clim. Past* **6**, 675–694 (2010).
17. Olson, D. M. *et al.* Terrestrial ecoregions of the world: a new map of life on earth. *BioScience* **51**, 933–938 (2001).
18. Ray, N. & Adams, J. M. A GIS-based vegetation map of the world at the last glacial maximum (25,000-15,000 BP). *Internet Archaeol.* **11**, (2001).
19. Takhtajan, A. L. *Floristic Regions of the World*. (University of California press, 1986).
20. Kreft, H. & Jetz, W. Global patterns and determinants of vascular plant diversity. *Proc. Natl. Acad. Sci. USA* **104**, 5925–5930 (2007).
21. Keil, P. & Chase, J. M. Global patterns and drivers of tree diversity integrated across a continuum of spatial grains. *Nat. Ecol. Evol.* **3**, 390–399 (2019).
22. Whittaker, R. H. *Communities and Ecosystems*. (Macmillan, 1975).
23. Körner, C. *et al.* A global inventory of mountains for bio-geographical applications. *Alp Botany* **127**, 1–15 (2017).

**Supplementary references 2 | References of Checklists and Floras from the Global Inventory of Floras and Traits (GIFT) used to compile the regional species composition data.**


1. Junak, S., Philbrick, R., Chaney, S. & Clark, R. *A checklist of vascular plants of Channel Islands National Park.* 2nd ed. (Southwest Parks and Monuments Association, Tucson, Arizona, 1997).

2. Broughton, D. A. & McAdam, J. H. A checklist of the native vascular flora of the Falkland Islands (Islas Malvinas). new information on the species present, their ecology, status and distribution. *The Journal of the Torrey Botanical Society* **132,** 115–148 (2005).

3. Kristinsson, H. *Checklist of the vascular plants of Iceland* (Náttúrufræðistofnun Íslands, Reykjavík, Iceland, 2008).

4. Sandbakk, B. E., Alsos, I. G., Arnesen, G. & Elven, R. The flora of Svalbard. Available at http://svalbardflora.no/ (1996).

5. Acevedo-Rodríguez, P. & Strong, M. T. Catalogue of the seed plants of the West Indies Website. Available at http://botany.si.edu/antilles/WestIndies/catalog.htm (2007).

6. Baker, M.L. & Duretto, M.F. *A census of the vascular plants of Tasmania* (Tasmanian Herbarium, Tasmanian Museum and Art Gallery, Hobart, Australia, 2011).

7. Tatewaki, M. Geobotanical studies on the Kurile Islands. *Acta Horti Gotoburgensis* **21,** 43–123 (1957).

8. Cheffings, C. M. & Farrell, L. (eds.). *The vascular plant red data list for Great Britain* (Joint Nature Conservation Committee, Peterborough, UK, 2005).

9. Stace, C.A., Ellis, R.G., Kent, D.H. & McCosh, D.J. *Vice-county Census Catalogue of the vascular plants of Great Britain, the Isle of Man and the Channel Islands* (Botanical Society of the British Isles, London, UK, 2003).

10. Conti, F., Abbate, G., Alessandrini, A. & Blasi, C. *Annotated Checklist of the Italian Vascular Flora* (Palombi Editori, Rome, Italy, 2005).

11. Case, T.J., Cody, M. L. & Ezcurra, E. *A new island biogeography of the Sea of Cortés* (Oxford University Press, New York, NY, 2002).

12. Shaw, J. D., Spear, D., Greve, M. & Chown, S. L. Taxonomic homogenization and differentiation across Southern Ocean Islands differ among insects and vascular plants. *J Biogeogr* **37,** 217–228 (2010).

13. Roux, J. P. *Synopsis of the Lycopodiophyta and Pteridophyta of Africa, Madagascar and neighbouring islands* (South African National Biodiversity Institute, Cape Town, South Africa, 2009).

14. 3D Environmental. Vegetation Communities and Regional Ecosystems of The Torres Strait Islands, Queensland, Australia. Torres Strait Regional Authority Land & Sea Management Unit, 2008.

15. 3D Environmental. Profile for management of the habitats and related ecological and cultural resource values of Mua Island. Torres Strait Regional Authority Land & Sea Management Unit, 2013.

16. Pelser, P. B., Barcelona, J. F. & Nickrent, D. L. Co's Digital Flora of the Philippines. Available at https://www.philippineplants.org/ (2011).

17. Barker, W. R., Barker, R. M., Jessop, J. P. & Vonow, H. P. Census of South Australian vascular plants. *Journal of the Adelaide Botanic Gardens Supplement* **1,** 1–396 (2005).

18. Belhacene, L. Catalogue 2010 des plantes vasculaires du département de la Haute-Garonne. *Supplément à Isaatis* **10,** 1–145 (2010).

19. BioScripts. Flora Vascular. Available at http://www.floravascular.com/ (2014).

20. Brennan, K. *An annotated checklist of the vascular plants of the Alligator Rivers Region, Northern Territory, Australia* (Supervising Scientist, Barton, Australia, 1996).

27. Funk, V. A., Hollowell, T., Berry, P., Kelloff, C. & Alexander, S. N. *Checklist of the plants of the Guiana Shield (Venezuela: Amazonas, Bolivar, Delta Amacuro; Guyana, Surinam, French Guiana)* (Department of Botany, National Museum of Natural History, Washington, DC, 2007).

28. Harris, D. J. *The vascular plants of the Dzanga-Sangha Reserve, Central African Republic* (Royal Botanic Garden Edinburgh, Edinburgh, Scotland, UK, 2002).

29. ZDSF & SKEW. Info Flora. Artenliste Schweiz 5x5 km. Available at https://www.infoflora.ch/de/daten-beziehen/artenliste-5x5-km.html (2014).

30. Zuloaga, F. O., Morrone, O. & Belgrano, M. Catálogo de las Plantas Vasculares del Cono Sur. Available at http://www.darwin.edu.ar/Proyectos/FloraArgentina/fa.htm (2014).

31. Kelloff, C. L. & Funk, V. A. *Preliminary checklist of the plants of Kaieteur National Park, Guyana* (National Museum of Natural History, Smithsonian Institution, Washington, 1998).

32. Le Houerou, H. N. Plant diversity in Marmarica (Libya & Egypt): a catalogue of the vascular plants reported with their biology, distribution, frequency, usage, economic potential, habitat and main ecological features, with an extensive bibliography. *Candollea* **59,** 259–308 (2004).

33. Lipkin, R. *Aniakchak National Monument and Preserve, vascular plant inventory: final technical report* (National Park Service, Southwest Alaska Network Inventory & Monitroing Program, Anchorage, USA, 2005).

34. Marticorena, C., Squeo, F. A., Arancio, G. & Muñoz, M. Catálogo de la flora vascular de la IV Región de Coquimbo. In *Libro rojo de la flora nativa y de los sitios prioritarios para su conservación: Región de Atacama,* edited by F. A. Squeo, G. Arancio & J. R. Gutiérrez (Ediciones Universidad de La Serena La Serena2008), pp. 105–142.

35. Nationalpark Eifel. Artenliste Farne und Blütenpflanzen. Available at http://www.nationalpark-eifel.de/go/artenliste.html (2015).

36. Norton, J. *et al. An illustrated checklist of the flora of Qatar* (Browndown Publications Gosport, Gosport, UK, 2009).

37. Pal, D., Kumar, A. & Dutt, B. Floristic diversity of Theog Forest Division, Himachal Pradesh, Western Himalaya. *Check list* **10,** 1083–1103 (2014).

38. Peña-Chocarro, M. d. C. *Updated checklist of vascular plants of the Mbaracayú Forest Nature Reserve (Reserva Natural del Bosque Mbaracayú), Paraguay* (Magnolia Press, Auckland, N.Z., 2010).

39. Pôle Flore Habitats. Catalogue de la flore vasculaire de Rhône-Alpes. Available at http://www.pifh.fr/pifhcms/index.php (2015).

40. Queensland Government. Census of the Queensland flora 2014. Available at https://data.qld.gov.au/dataset/census-of-the-queensland-flora-2014 (2014).

41. Rundel, P. W., Dillon, M. O. & Palma, B. Flora and Vegetation of Pan de Azúcur National Park in the Atacama desert of Northern Chile. *Gayana Bot* **53,** 295–315 (1996).

42. SLUFG. *Rote Liste und Artenliste Sachsen. Farn- und Samenpflanzen* (Sächsisches Landesamt für Umwelt, Landwirtschaft und Geologie, Dresden, Germany, 2015).

43. SANBI. Plants of Southern Africa. An online checklist. Available at http://posa.sanbi.org (2014).

44. Selvi, F. A critical checklist of the vascular flora of Tuscan Maremma (Grosseto province, Italy). *Fl. Medit* **20,** 47–139 (2010).

45. Short, P. S., Albrecht, D. E., Cowie, I. D., Lewis, D. L. & Stuckey, B. M. *Checklist of the vascular plants of the Northern Territory* (Department of Natural Resources, Environment, The Arts and Sport, Darwin, 2011).

46. USDA & NRCS. The PLANTS Database. Available at http://plants.usda.gov (2015).

47. Viciani, D., Gonnelli, V., Sirotti, M. & Agostini, N. An annotated check-list of the vascular flora of the "Parco Nazionale delle Foreste Casentinesi, Monte Falterona e Campigna"(Northern Apennines Central Italy). *Webbia* **65,** 3–131 (2010).

48. Vogt, C. Composición de la Flora Vascular del Chaco Boreal, Paraguay. I. Pteridophyta y Monocotiledoneae. *Steviana* **3,** 13–47 (2011).

49. van Vreeswyk, A.M.E., Payne, A.L., Leighton, K. A. & Hennig, P. An inventory and condition survey of the Pilbara region, Western Australia. Department of Agriculture, Government of Western Australia, 2004.

50. Stalmans, M. *Tinley's plant species list for the Greater Gorongosa ecosystem, Moçambique.* Unpublished report by International Conservation Services to the Carr Foundation and the Ministry of Tourism (2006).

51. VicFlora. Flora of Victoria. Available at http://vicflora.rbg.vic.gov.au (2016).

52. Oggero, A. J. & Arana, M. D. Inventario de las plantas vasculares del sur de la zona serrana de Córdoba, Argentina. *Hoehnea* **39,** 171–199 (2012).

53. Masharabu, T. Flore et végétation du Parc National de la Ruvubu au Burundi: diversité, structure et implications pour la conservation. PhD Thesis. Université Libre de Bruxelles, 2011.

59. Евстигнеев, О. И. & Федотов, Ю. П. *Флора сосудистых растений заповедника "Брянский лес"* (Гос. природ. биосфер. заповедник Брян. лес, Брянск, 2007).

60. Lazkov, G. A. & Sultanova, B. A. *Checklist of vascular plants of Kyrgyzstan* (Botanical Museum, Finnish Museum of Natural History, Helsinki, 2011).

61. Pandža, M. Flora parka prirode Papuk (Slavonija, Hrvatska). *Šumarski list* **134,** 25–43 (2010).

62. Tropicos. Catálogo de las Plantas Vasculares de Bolivia. Available at http://www.tropicos.org/Project/BC (2015).

63. Tropicos. Catalogue of the Vascular Plants of Ecuador. Available at http://www.tropicos.org/Project/CE (2015).

64. Tropicos. Panama Checklist. Available at http://www.tropicos.org/Project/PAC (2015).

65. Zvyagintseva, K. O. *An annotated checklist of the urban flora of Kharkiv* (Kharkiv National University, Kharkiv, Ukraine, 2015).

66. Notov, A. A. *National park "Zavidovo": vascular plants, bryophyte, lichens* (Delovoj mir, Moscow, 2010).

67. Хапугин, А. А. *Сосудистые растения Ромодановского района Республики Мордовия (конспект флоры)* (Saransk, 2013).

68. Rakov, N. S., Saksonov S.V., Senator S.A. & Vasjukov V.M. *Vascular plants of Ulyanovsk Region* (Russian Academy of Sciences, Togliatti, Russia, 2014).

69. Conti, F. & Bartolucci, F. *The Vascular Flora of the National Park of Abruzzo, Lazio and Molise (Central Italy)* (Springer, Barisciano, Italy, 2015).

70. Botanical Garden Tel Aviv, personal communication.

81. Kupriyanov, A. N. & Kupriyanov, O. A., personal communication.

82. Revushkin, A. *Key to the flora of the Tomsk Region [Opredelitel rasteniy Tomskoy oblasty].* with minor corrections by Alexandr Ebel (Tomsk State University Press, Tomsk, Russia, 2014).

83. Silantyeva, M. *Synopsis of the flora of Altayskyi Krai [Konspekt flory Altayskogo kraya].* with minor corrections by Alexandr Ebel (Altay State University Press, Barnaul, Russia, 2013).

84. Thomas, J. Plant diversity of Saudi Arabia. Flora checklist. Available at http://plantdiversityofsaudiarabia.info/biodiversity-saudi-arabia/flora/Checklist/Cheklist.htm (2011).

85. Krestov, P. V., personal communication.

86. Lorite, J. An updated checklist of the vascular flora of Sierra Nevade (SE Spain). *Phytotaxa* **261,** 1–57 (2016).

87. Nowak, A., personal communication.

88. Ministry of Environment. *The national red list 2012 of Sri Lanka. Conservation status of the fauna and flora* (Colombo, Sri Lanka, 2012).

89. Silaeva, T. B. *et al.* Flora of the national park "Smolny". Mosses and vascular plants: annotated list of species. Commission of RAS for the Conservation of Biological Diversity, 2011.

90. Karlsson, T. & Agestam, M. Checklist of Nordic vascular plants. Sweden Checklist. Available at http://www.euphrasia.nu/checklista/index.eng.html (2014).

91. Dimopoulos, P. *et al. Vascular plants of Greece. An annotated checklist* (Botanischer Garten und Botanisches Museum Berlin-Dahlem, Freie Universität Berlin; Hellenic Botanical Society, Berlin, Athens, 2013).

92. Bundesamt für Naturschutz. Floraweb. Available at www.floraweb.de (2016).

93. Cochard, R. & Bloesch, U. Electronic plant species database of the Saadani National Park, coastal Tanzania. Available at http://www.wildlife-baldus.com/saadani.html (2007).

94. Hyde, M. A., Wursten, B. T., Ballings, P. & Coates Palgrave, M. Flora of Botswana. Available at http://www.botswanaflora.com/ (2016).

95. Bingham, M. G., Willemen, A., Wursten, B. T., Ballings, P. & Hyde, M. A. Flora of Zambia. Available at http://www.zambiaflora.com/ (2016).

96. Hyde, M. A., Wursten, B. T., Ballings, P. & Coates Palgrave, M. Flora of Zimbabwe (2016).

97. Hyde, M. A., Wursten, B. T., Ballings, P. & Coates Palgrave, M. Flora of Mozambique. Available at http://www.mozambiqueflora.com/ (2016).

98. Hyde, M. A., Wursten, B. T., Ballings, P. & Coates Palgrave, M. Flora of Malawi. Available at http://www.malawiflora.com/index.php (2016).

99. Fischer, M. A., Adler, W. & Oswald, K. *Exkursionsflora für Österreich, Liechtenstein und Südtirol. Bestimmungsbuch für alle in der Republik Österreich, im Fürstentum Liechtenstein und in der Autonomen Provinz Bozen.* 3rd ed. (Oberöstereichisches Landesmuseum, Linz, Austria, 2008).

100. Wieringa, J. J., personal communication.

101. Chang, C.-S., Kim, H. & Chang, K. *Provisional Checklist of the Vascular Plants for the Korea Peninsular Flora (KPF). Version 1.0* (Korea, 2014).

108. Turner, I. M. *A catalogue of the vascular plants of Malaya* (Gardens' Bulletin, Singapore, 1995).

109. Western Australian Herbarium. FloraBase - the Western Australian Flora. Available at https://florabase.dpaw.wa.gov.au/ (2017).

110. Royal Botanic Gardens and Domain Trust. PlantNET - The NSW Plant Information Network System. Available at http://plantnet.rbgsyd.nsw.gov.au (2017).

111. Breckle, S.-W., Hedge, I. C. & Rafiqpoor, M. D. *Vascular plants of Afghanistan. An augmented checklist* (Scientia Bonnensis, Bonn, 2013).

112. Landcare Research New Zealand. Ecological Traits of New Zealand Flora. Available at https://ecotraits.landcareresearch.co.nz/.

113. Yonekura, K. & Kajita, T. BG Plants Japanese name and Scientific name Index (YList). Available at http://ylist.info (2013).

114. Backer, C. A. & Bakhuizen van den Brink, RC. *Flora of Java.* (Spermatophytes only) (Noordhoff, Groningen, 1963).

115. Euro+Med. Euro+Med PlantBase - the information resource for Euro-Mediterranean plant diversity. Available at http://ww2.bgbm.org/EuroPlusMed/ (2006).

116. Ulloa Ulloa, C. *et al.* An integrated assessment of the vascular plant species of the Americas. *Science (New York, N.Y.)* **358,** 1614–1617; 10.1126/science.aao0398 (2017).

117. Yena, A. V. *Prirodnaja flora krymskogo poluostrova. Spontaneous flora of the Crimean Peninsula* (Orianda, Simferopol', 2012).

118. Tropicos. Catalogue of the Vascular Plants of Madagascar. Available at http://www.tropicos.org/Project/MADA (2020).

119. Newman, M. *A checklist of the vascular plants of Lao PDR* (Royal Botanic Garden Edinburgh, Edinburgh, Scotland, UK, 2007).

120. Kluge, J. *et al.* Elevational seed plants richness patterns in Bhutan, Eastern Himalaya. *J. Biogeogr.* **44,** 1711–1722; 10.1111/jbi.12955 (2017).

121. Patzelt, A., personal communication.

122. Rodriguez, R. *et al.* Catálogo de las plantas vasculares de Chile. *Gayana Bot.* **75,** 1–430; 10.4067/S0717-66432018000100001 (2018).

123. Hutchinson, J., Dalziel, J. M., Keay, R.W.J. & Hepper, N. *Flora of West Tropical Africa* (Crown agents for overseas governments, London, UK, 2014).

124. Cámara-Leret, R. *et al.* New Guinea has the world's richest island flora. *Nature* **584,** 579–583; 10.1038/s41586-020-2549-5 (2020).

125. Elizabeth Joyce *et al.* Checklist of the vascular flora of the Sunda-Sahul Convergence Zone. *Pensoft Publishers* **8,** e51094; 10.3897/BDJ.8.e51094 (2020).

126. BGCI. GlobalTreeSearch online database (version 1.4). Available at https://tools.bgci.org/global_tree_search.php (2021).

127. Inderjit, personal communication.

128. Hassler, M. World Ferns. Synonymic Checklist and Distribution of Ferns and Lycophytes of the World. Available at www.worldplants.de/ferns/ (2020).
